# Immune memory shapes human polyclonal antibody responses to H2N2 vaccination

**DOI:** 10.1101/2023.08.23.554525

**Authors:** Yuhe R. Yang, Julianna Han, Hailee R. Perrett, Sara T. Richey, Abigail M. Jackson, Alesandra J. Rodriguez, Rebecca A. Gillespie, Sarah O’Connell, Julie E. Raab, Lauren Y. Cominsky, Ankita Chopde, Masaru Kanekiyo, Katherine V. Houser, Grace L. Chen, Adrian B. McDermott, Sarah F. Andrews, Andrew B. Ward

**Author notes:** Correspondence to, ORCID 0000-0001-7153-3769;, ORCID 0000-0002-2583-1949. These authors contributed equally: Yuhe R. Yang, Julianna Han, Hailee R. Perrett, and Sarah F. Andrews.

## Abstract

Influenza A virus subtype H2N2, which caused the 1957 influenza pandemic, remains a global threat. A recent phase I clinical trial investigating a ferritin nanoparticle displaying H2 hemagglutinin in H2-naïve and H2-exposed adults. Therefore, we could perform comprehensive structural and biochemical characterization of immune memory on the breadth and diversity of the polyclonal serum antibody response elicited after H2 vaccination. We temporally map the epitopes targeted by serum antibodies after first and second vaccinations and show previous H2 exposure results in higher responses to the variable head domain of hemagglutinin while initial responses in H2-naïve participants are dominated by antibodies targeting conserved epitopes. We use cryo-EM and monoclonal B cell isolation to describe the molecular details of cross-reactive antibodies targeting conserved epitopes on the hemagglutinin head including the receptor binding site and a new site of vulnerability deemed the medial junction. Our findings accentuate the impact of pre-existing influenza exposure on serum antibody responses.

**Highlights:**

- Serum Abs after first H2-F vaccination in H2-exposed donors bound variable HA head epitopes
- Serum Abs after first H2-F vaccination in H2-naïve donors bound conserved HA head and stem epitopes
- RBS-targeting VH1-69 cross-reactive antibodies were induced in H2-naïve individuals
- The medial junction is a previously uncharacterized conserved epitope on the HA head

## Introduction

Responsible for causing five pandemics within the past 110 years alone, influenza viruses are one of the greatest threats to mankind. During non-pandemic years, influenza-related complications affect millions of people^1^ https://www.cdc.gov/flu/about/burden/index.html, impacting their daily lives and the global economy. Soberingly, pandemic influenza is a constant threat as the virus can undergo antigenic shift within the vast animal reservoir and cross the species barrier^2^, such as what occurred during the 1918 Spanish flu pandemic which resulted in 50–100 million deaths. Pandemic influenza often features surface glycoproteins for which the human population is largely or wholly naïve^3,4^, necessitating a more thorough understanding of the immune recognition of influenza subtypes by the general populace to better inform disease surveillance and pandemic prediction efforts.

Influenza A viruses are categorized by their surface glycoproteins including hemagglutinin (HA), which binds sialic acid receptors on the surface of a host cell and mediates fusion of the virus with the host endosomal membrane. Humoral immune responses to HA are known to be protective against infection by influenza^5,6^ and are readily induced post-infection or post-vaccination. Yet the success of these antibody responses—along with additional factors—drives the influenza virus to accumulate mutations to adapt its HA^7^, a process known as antigenic drift.

To prevent a future pandemic, eliciting broadly protective immunity through vaccination is our best line of defense. Antibody responses to the HA head domain are immunodominant^8^ and often highly effective in neutralizing specific viral strains.^9^ The head domain is highly susceptible to antigenic drift^10–12^ enabling the virus to escape these responses. Antibodies elicited by infection or vaccination that target conserved sites on HA—such as in the stem domain^13–17^—can offer protection through direct neutralization of the virus or through recruitment of adaptive and innate immune defenses to sites of infection. Various vaccination strategies, such as using novel influenza virus strains, chimeric HAs, and mosaic HAs, have shown promise in generating broadly cross-reactive and protective antibodies to these sites.^18–20^

A truly universal influenza vaccine must generate broadly neutralizing responses against the 18 recognized HA subtypes—especially those implicated in recent human pandemics: H1, H2, and H3. While only the H1N1 and H3N2 influenza A subtypes currently circulate in humans, the H2N2 influenza virus poses a distinct risk to humans. H2N2 was the causative agent of the 1957 ‘Asian flu’ pandemic, which originally emerged from an avian reservoir.^21^ This subtype resulted in more than 1 million deaths and circulated among the human population from 1957 until 1968 before being replaced by the H3N2 subtype. Yet, H2-influenza viruses continue to infect farm animals, birds, and swine.^22–24^ Further, the H2N2 HA sequence is highly conserved between human and avian species, resulting in an ever-present risk of interspecies transmission of H2N2 and the potential to trigger a new influenza pandemic, especially considering H2-specific immunity in humans exposed to H2N2 viruses pre-1968 has been waning.^25^ Thus, comprehensive analysis of human antibody responses and how these responses differ between age groups is crucial to gauge the effectiveness of candidate influenza virus vaccines. People born before 1968 likely have pre-existing immunity to H2N2 viruses due to childhood exposure. Conversely, younger populations born after 1968 are naïve to the H2N2 subtype, having only been exposed to seasonal H1N1 and H3N2 strains.^26^ These populations represent an excellent cohort to assess the vaccination strategy of expanding pre-existing antibody responses from one subtype (in this case, H1N1) to the conserved sites of another (H2N2).

A recent human phase I clinical trial (NCT03186781) assessed this vaccination strategy using a H2 HA ferritin nanoparticle (H2-F) as the antigen.^27^ Previous characterization of responses to the H2 antigen in this trial indicated that H2-naïve individuals generated cross-reactive serological and B-cell responses to the H2 stem. ^27,28^ Those with pre-existing immunity demonstrated more H2-specific serological antibody responses not targeting the H2 stem.^27,28^ These results align with our previous work using electron microscopy polyclonal epitope mapping (EMPEM) to demonstrate at the structural level that novel vaccination biases initial immune responses to conserved sites in novel and seasonal influenza vaccinations.^9,29–31^

In this work, we use EMPEM and complementary serological analyses to assess polyclonal antibody (pAb) responses in different age cohorts and observe that initial exposure to H2 HA generates cross-reactive pAb responses to the receptor binding site (RBS). Conversely, secondary exposure generates diverse, strain-specific responses. We found that the H2-naïve individuals likely recalled cross-reactive pAb responses from pre-existing immunity to H1N1 viruses. The molecular details of cross-reactive and strain-specific monoclonal antibodies (mAbs) isolated from H2-F-vaccinated individuals were revealed by high resolution cryo-electron microscopy (cryo-EM). We also describe a broadly cross-reactive antibody to a previously unappreciated epitope on HA containing conserved residues in the central helix of HA2 and the vestigial esterase domain. This new ‘medial junction’ epitope likely adds an additional layer of protection against diverse influenza viruses. Overall, by characterizing recalled homo- and hetero-subtypic pre-existing immunity after H2-F vaccination, this study adds critical immunological knowledge for optimizing influenza vaccines.

## RESULTS

### Vaccine-induced antibody responses in naïve and pre-exposed individuals

A recent human phase I clinical trial (NCT03186781) investigated the safety and immunogenicity of two experimental H2N2-based (A/Singapore/1/1957) influenza vaccines: (1) VRC-FLUDNA082-00-VP, a plasmid DNA vaccine encoding full-length influenza A H2 and (2) VRC-FLUNPF081-00-VP, a ferritin nanoparticle presenting multivalent H2 ectodomains.^27,32^ To test safety and immunogenicity of these experimental vaccines, fifty human participants were vaccinated with different prime/boost strategies.^27^ As humans can be naïve or have pre-existing immunity to H2N2 viruses, the participants were divided into two age groups—those born after 1968 without pre-existing H2N2 immunity and those living before H2N2 viruses ended circulation in 1968 (Fig. 1A, groups 1**–**2 and groups 3**–**4, respectively). In both age cohorts, one group received a primary vaccination of H2 DNA plasmid antigen followed by a secondary vaccination with H2-F while the other group received a primary and secondary vaccine regimen of H2-F vaccine (Fig. 1A, groups 1 & 3 and groups 2 & 4, respectively). Serum samples from 12 representative participants, 3 from each group, were collected at weeks 0, 4, 16, and 20 (week 4 post-boost; Fig. 1B). Using a Meso Scale Discovery (MSD) assay, we observed increases in H2 HA-specific serum antibody titers over the course of vaccination in all 12 participants (Fig. 1C).^27^

**Fig. 1.**
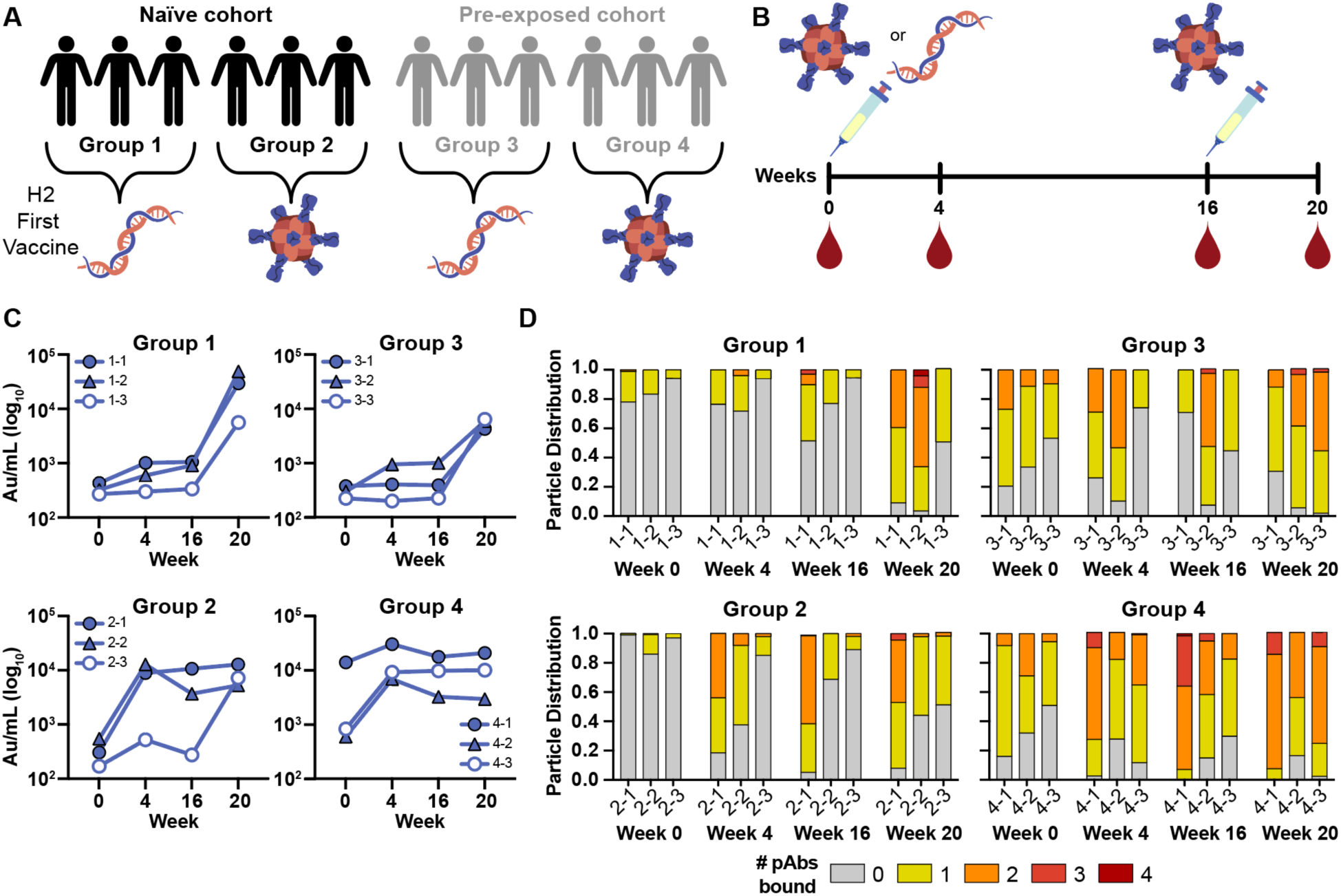
H2N2 vaccine elicits antigen-specific immune responses in trial participants. (A) Schematic of the H2N2 vaccine trial. Participants were placed into four groups separated by exposure status and vaccination platform: naïve participants who were primed with H2 DNA plasmid-based vaccine (group 1) or the multivalent H2-F nanoparticle (group 2), and pre-exposed participants who were first vaccinated with the DNA plasmid-based vaccine (group 3) or the H2-F nanoparticle (group 4). All groups received secondary vaccinations with H2-F. Individual participants are notated by -1, -2, and -3 for a total of n=3 per group. (B) DNA plasmid and H2-F antigens were administered in two immunizations: first vaccine dose at week 0 and second vaccine dose at week 16. Serum samples were collected at weeks 0, 4, and 16 (after the first vaccination) and week 20 (after boost). (C) MSD binding levels of serum antibodies against H2 HA ectodomain of human participants as measured using Au/ml, arbitrary units/ml. (D) nsEMPEM semi-quantitative epitope occupancy analysis denoting the proportion of HA trimers with 0, 1, 2, 3, or 4 pAbs bound (grey, yellow, orange, dark orange, and red, respectively) for each participant, noted on the x-axis.

To further probe the dynamics of serum pAb responses to H2 in this trial, we used negative stain EMPEM (nsEMPEM)^30,31^ to map pAb-targeted epitopes on homologous H2 HA (A/Singapore/1/1957). Participant pAbs were digested to Fab, complexed in excess with HA, purified, and imaged. We analyzed the proportion of particles in each 2D class based on the number of pAbs bound to HA. Consistent with sera antibody binding to immobilized H2 HA (Fig. 1C), the epitope occupancy increased as more pAbs were elicited by vaccination, highlighting the immunogenicity of the H2 vaccines (Fig. 1D). While the two groups who received the DNA primary vaccination showed a moderate increase in antibody titers and epitope occupancy after the priming immunization, all groups showed sustained and increased pAb responses after H2-F primary and/or secondary vaccination (Fig. 1C and D).

### Characterizing pAb responses to head and stem epitopes on H2 HA

Previous studies of antibody responses to influenza infection and vaccination revealed cross-reactive HA stem responses are usually recalled upon exposure to antigenically novel influenza strains while strain-specific head responses are recalled upon re-exposure to strains encountered previously.^8,30,33,34^ As these differential responses have major implications for strain-specific and cross-reactive immunity, we generated epitope landscapes of pAb responses at each time point by nsEMPEM (Fig. 2; Fig. S1 and S2; Table S1). We observed clear differences in pre-existing immunity to homologous H2 HA for naïve and pre-exposed groups at the serum level (Fig. 2A and C). For the six H2-naïve human participants, only stem-specific pAbs were observed at week 0, which were likely pre-existing, cross-reactive antibodies elicited by prior exposure to seasonal influenza infection or vaccination. After the first vaccination dose, pAb responses expanded to target the RBS. Responses further diversified to target variable head and vestigial esterase epitopes after the H2-F boost (Fig. 2A and B). In contrast, the majority of pre-exposed participants demonstrated baseline pAbs targeting head and stem epitopes, which expanded after the H2-F first and or second vaccine dose to target variable head epitopes (Fig. 2C and D). Together, these data demonstrate both vaccination strategies induce strong pAb responses targeting epitopes with higher sequence conservation upon primary exposure to H2 HA, while secondary exposure to H2 HA induces more diversified immunity to variable epitopes on the HA head.

**Fig. 2.**
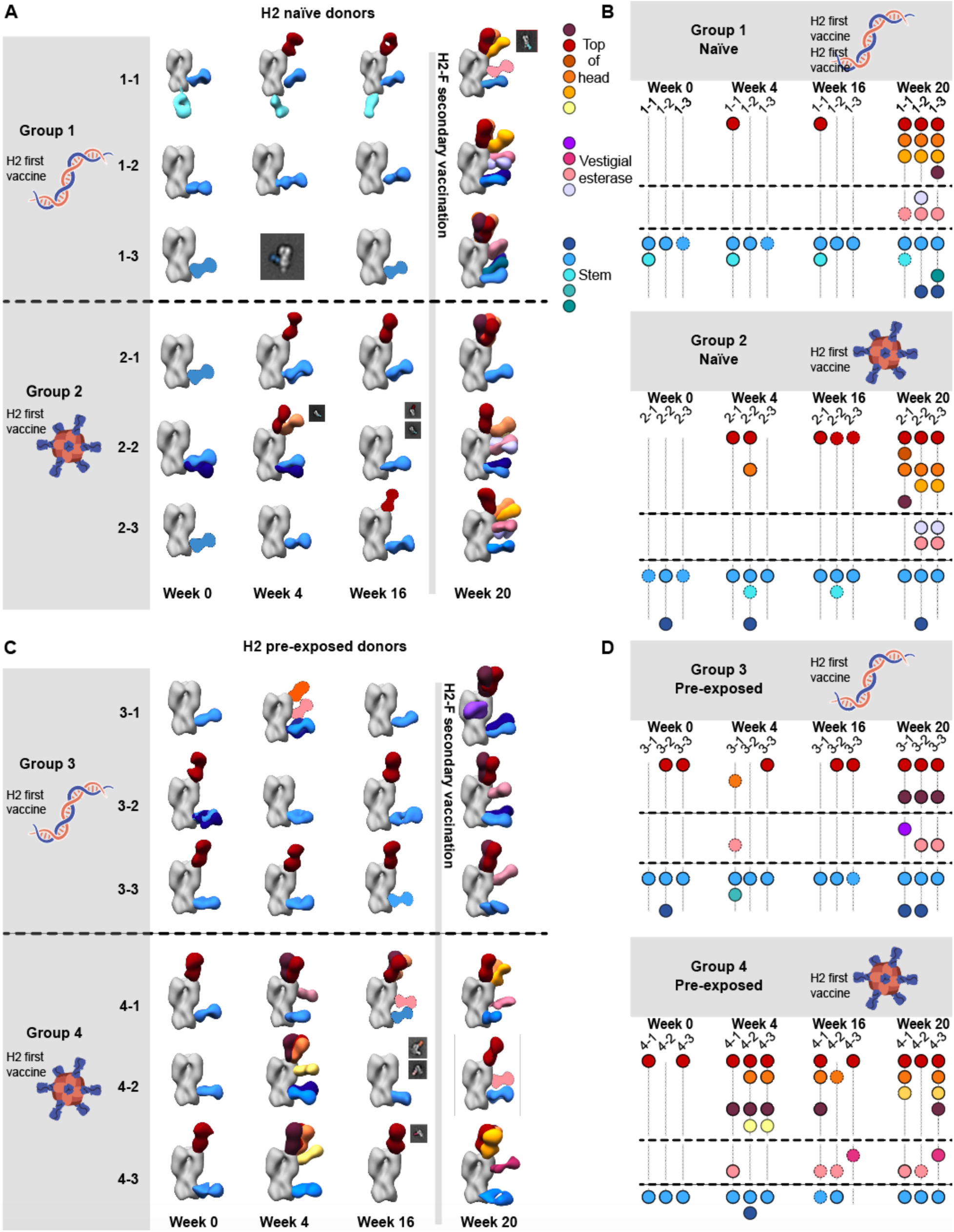
Polyclonal analysis of H2N2 vaccine trial participants. (A & C) Composite 3D reconstructions with segmented pAb specificities of each participant displayed on one protomer of the H2 HA trimer (grey) for naïve participants (A) or pre-exposed participants (C). Gray lines indicates whether samples were collected pre or post-H2F boost at week 16. Fabs represented as 2D class averages or depicted on the H2 HA trimer as a silhouette with dotted outline have limited particle representation and/or low particle abundance, and their epitopes were consequently predicted. Epitope cluster color scheme is shown on the right. (B & D) Summary of pAb specificities for each group. Each circle represents a unique pAb specificity denoted by the color scheme in A & C.

While pAb maps of each timepoint demonstrated the diversity of epitopes targeted before and after vaccination, we hypothesized there would be clear differences in the frequency of head versus stem responses. Thus, we analyzed overall trends of epitope distribution between serum samples using semi-quantitative EMPEM and MSD analyses (Fig. 3). 3D sorting and particle counting enabled semi-quantitative nsEMPEM analysis of the distribution of head- or stem-specific immune complexes (Fig. 3A). In tandem, we performed plate-based sera binding analyses using full-length H2 HA or H2 HA stem to assess a participant’s relative ratio of head and stem-targeting pAbs (Fig. 3B). Both methods converged on similar trends: stem-targeting pAbs were more prominent at week 0 while head-specific antibodies dominated the pAb landscape at week 20, prompted by the H2-F boost at week 16. Overall, primary H2 vaccination with the H2-F nanoparticle boosted pAb responses to the stem while secondary exposure, whether through the second immunization in the naïve groups or through the first immunization in pre-exposed participants, elicited diverse pAb responses to head epitopes on H2 HA

**Fig. 3.**
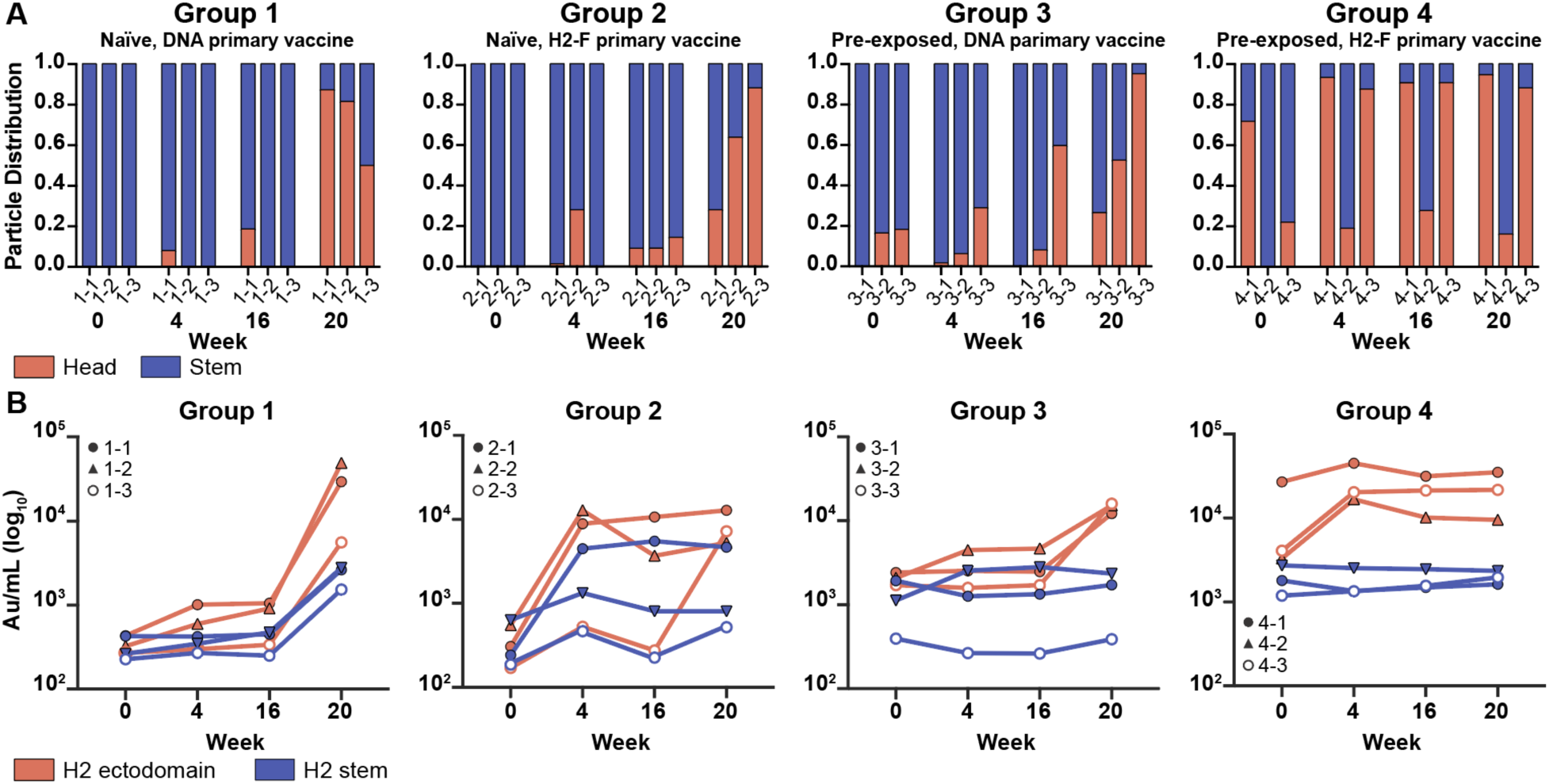
Frequency of H2 HA head and stem responses. (A) nsEMPEM semi-quantitative H2 HA epitope occupancy analysis indicating the proportion of pAb-containing particles in 2D classes targeting the head (orange) or stem (blue). (B) Serum antibody titers measured by MSD using probes of HA ectodomain (orange) and HA stem (blue). Serum samples of each participant are presented with unique symbols.

### Cross-reactivity of vaccine-induced head targeting antibodies to H1

To further explore immunity elicited by H2 vaccination, we tested the ability of serum pAbs to interact with H1 HA by nsEMPEM. Evaluating H2 naïve participants 1-1 and 2-2, we observed diverse pre-existing immunity to H1 with pAbs targeting epitopes on the head and stem (Fig. 4A). Both donors had pre-existing immunity to the H2 HA stem epitopes and upon vaccination elicited responses to the HA head (Fig. 4A). Neither participant had H2 head-targeting pAb responses at week 0, but after H2 vaccination both participants elicited responses to the RBS of H2. In summary, pre-existing stem responses most likely elicited from prior H1 exposure can cross-react with H2 HA while H2 HA vaccination further elicited strain-specific pAbs to the RBS.

**Fig. 4.**
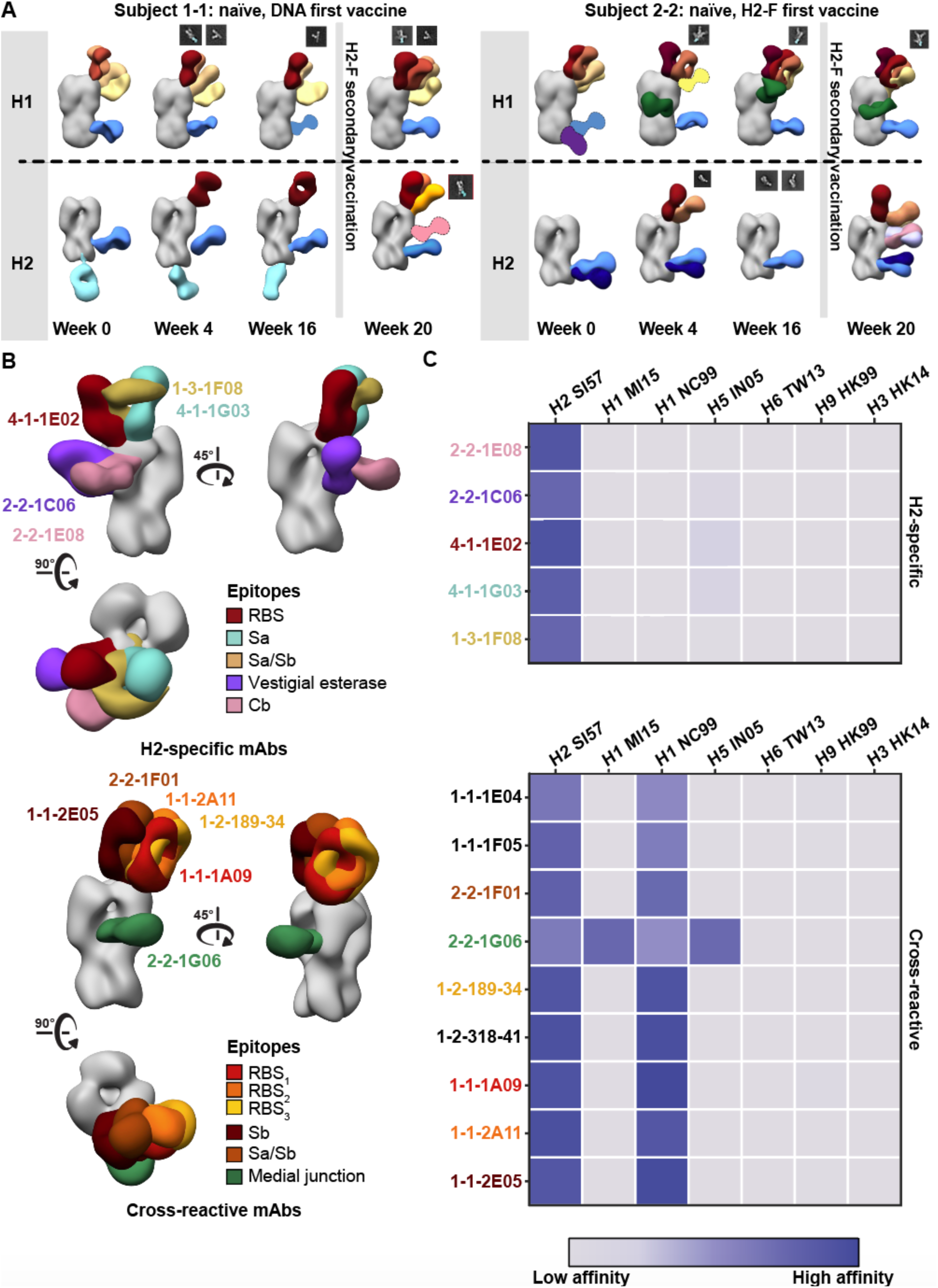
Cross-reactivity of elicited immune responses. (A) Segmented nsEM 3D reconstructions of participant 1-1 (left) and 2-2 (right) pAbs complexed with either H1/NC99 or H2/1957 HA antigen. Fabs represented as 2D class averages or depicted on the H2 HA trimer as a silhouette with dotted outline have limited particle representation and/or low particle abundance, and their epitopes were consequently predicted. Gray lines indicates whether samples were collected pre- or post-H2F secondary vaccination at week 16. (B) Representative nsEM reconstructions of H2-specific (top) and H1-cross reactive (bottom) monoclonal antibodies in complex with H2 HA. (C) Binding levels of mAbs isolated from plasmablasts or memory B cells against HA subtypes 1 and 2 weeks after H2-F boost.

To assess epitopes targeted by cross-reactive and H2-specific head-binding antibodies, mAbs from vaccine-elicited B cells were isolated and characterized from six participants from blood collected 1 and 2 weeks after H2-F boost (Fig. 4B and C, Table S3). nsEM epitope mapping revealed distinct patterns for each mAb group: H2-specific mAbs targeted a variety of epitopes on the HA head including RBS, vestigial esterase, and antigenic sites Sa and Cb while the majority of cross-reactive mAbs targeted the RBS (Fig. 4B). These cross-reactive RBS antibodies were detected in three subjects (1-1, 1-3, and 2-2), demonstrating this broad antibody response is seen across individuals after H2 vaccination.

We next applied high-resolution cryoEMPEM to a previously naïve participant (1-1, week 20) to see if pAbs resembling the cross-reactive mAbs isolated from B cells in the same individual could be detected. Consistent with nsEMPEM findings (Fig. 2A, Fig. S3A and B), cryoEMPEM analysis of participant 1-1 revealed RBS- and stem-targeting pAbs (Fig. 5A and Fig. S3C and D). We obtained two high-resolution reconstructions corresponding to unique pAb complexes that targeted the RBS with distinct angles of approach (Fig. 5A; Table S2, Fig. S4 and S5). We found the pAbs overlapped with low-resolution maps of mAbs 1-1-2E05 and 1-1-1F05 isolated from participant 1-1’s B-cells 1**–**2 weeks after the H2-F boost. This suggests these mAbs represent the corresponding antibody response within each pAb specificity (Fig. 5A). For a direct comparison, we solved the structure of 1-1-1F05, which represents an abundant head-specific lineage within the B-cell population (Fig. 5B), in complex with H2 HA at high-resolution using cryoEM (Fig 5C, top). Remarkably, 1-1-1F05 demonstrated high structural similarity with pAb_2 (Fig. 5C, bottom). In the model of 1-1-1F05, we observed the CDRH3 loop’s interactions at the RBS epitope were in high agreement with the density map of pAb_2, heavily suggesting 1-1-1F05’s cross-reactivity is indeed a large component of the circulating serum response to the RBS.

**Fig. 5.**
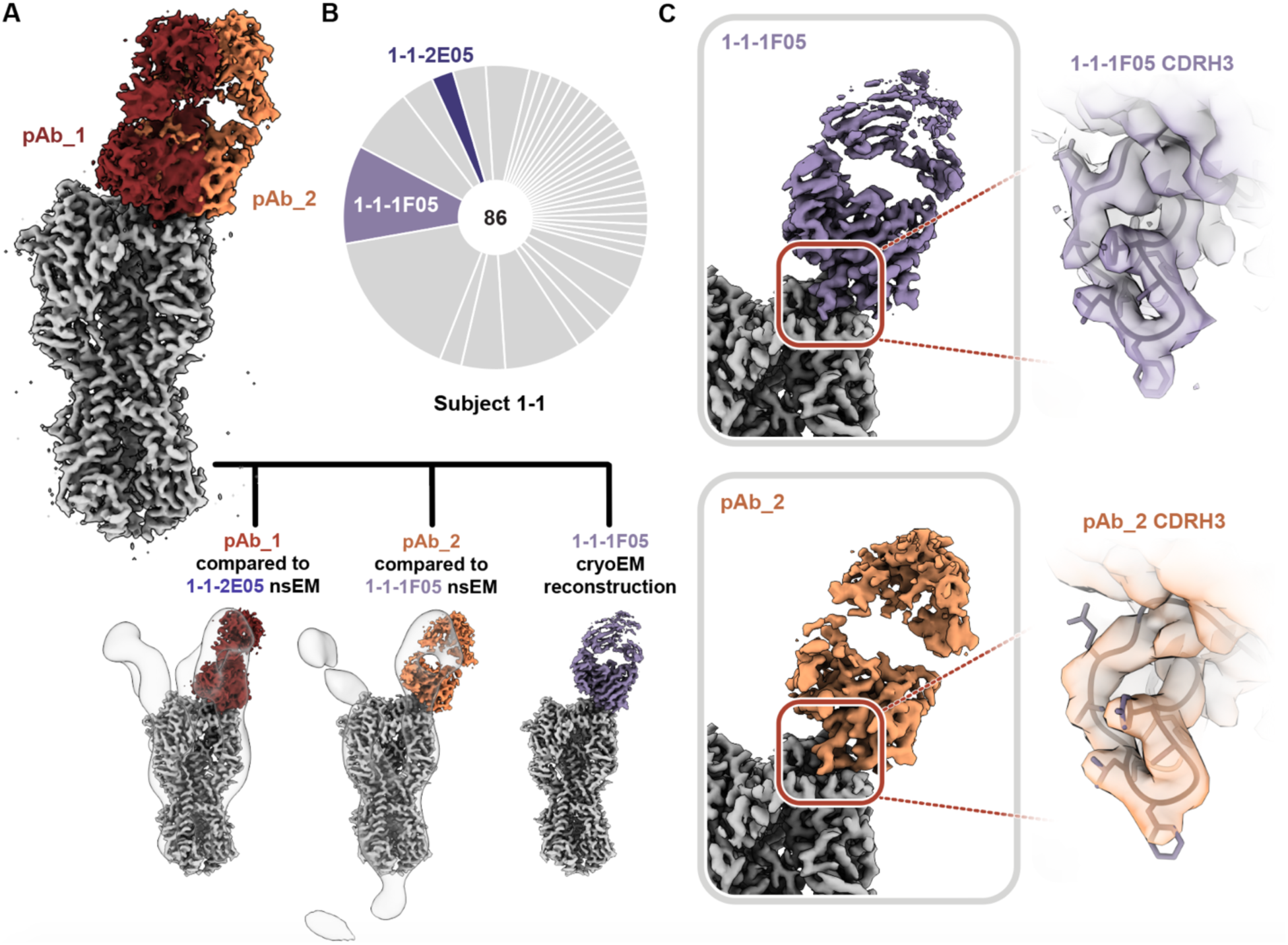
Structural analysis of RBS-targeting pAbs in participant 1-1. (A) CryoEMPEM analysis of immune complexes from participant 1-1 on week 20. H2 HA antigen is colored grey with two segmented Fab density maps colored in red and orange (top). nsEM maps of monoclonal antibodies in complex with H2 HA are overlaid against the corresponding cryo-EM map (bottom). (B) Pie chart showing Ig repertoire of single-cell sorted and sequenced H2-head specific plasmablasts from participant 1-1 one week after the H2HA Ferritin boost. (C) Single-particle cryo-EM reconstruction of H2 HA in complex with 1-1-1F05. (D) Density maps at the epitope-paratope interaction of 1-1-1F05 (top) and pAb_2 (bottom). The atomic model of 1-1-1F05 is shown in purple and docked into both density maps.

### Structural analysis of isolated mAbs describes binding mechanism similar to known RBS-targeting mAbs

To further dissect the differences in head targeting-antibodies (Fig. 4 and 5), we generated a high-resolution map of H1 cross-reactive RBS-targeting mAb 1-1-1E04 in addition to 1-1-1F05. Additionally, we investigated two H2-specific mAbs that target the RBS and antigenic site Sa (4-1-1E02 and 4-1-1G03, respectively; Fig. 6A and Fig. S6). For the cross-reactive mAbs 1-1-1F05 and 1-1-1E04, we also define their interactions with H1 (strain A/New Caledonia/20/1999(H1N1); NC99; Fig. S7). All maps were of sufficient quality to build atomic models of HA bound to Fab except in the case of H2’s cleavage site, which had heterogeneous density and was omitted from H2 models.

**Fig. 6.**
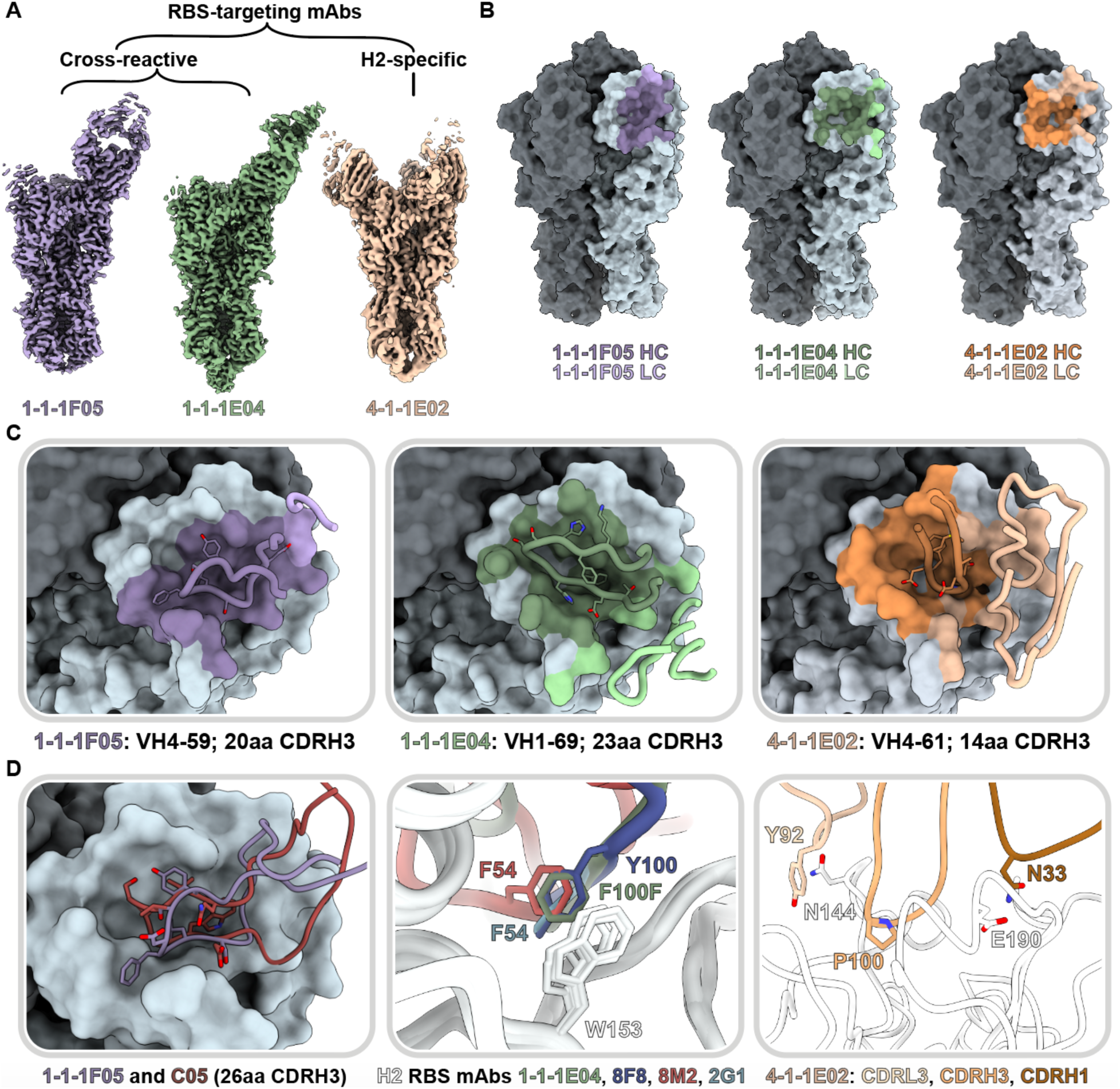
Structural characterization of RBS-targeting antibodies. (A) CryoEM density maps of mAb-HA complexes. (B) Antibody footprints of 1-1-1F05, 1-1-1E04, and 4-1-1E02 mAbs on HA colored to indicate heavy and light chain interactions. (C) Antibody loop interactions with the RBS pocket, with key CDRH3 residues shown. CDRH3 residue lengths are annotated using the IMGT numbering scheme. (D) 1-1-1F05 and bnAb C05 (PDB 4FP8) CDRH3 loops superimposed (left); 1-1-1E04 superimposed with bnAbs 2G1 (PDB 4HF5) and 8M2 (PDB 4HFU) and 8F8 (PDB 4HF5, middle); and 4-1-1G03 epitope-paratope interaction with key side chains shown (right). (E) Sequence alignment of CDRH3 loops shown in descending order by length.

Both cross-reactive mAbs 1-1-1F05 and 1-1-1E04 target the RBS from a single angle of approach. Relative to one another, there is an approximately 90° rotation in the heavy and light chains (Fig. 6A). The CDRH3 loops of these mAbs—20 residues for 1-1-1F05 and 23 for 1-1-1E04—insert into the RBS and mediate the majority of interactions with their epitopes (Fig. 6B and C). In contrast, strain-specific mAb 4-1-1E02 had a larger epitope footprint encompassing the RBS. Its heavy and light chains mediate its interactions at the HA surface, which includes contributions from a 14-residue long CDRH3 loop (Fig. 6B and C). The non-RBS binding mAb 4-1-1G03 targets antigenic site Sa interacting across two protomers with contributions from its light and heavy chains (Fig S6).

As the RBS is functionally conserved, antibodies to this site have the potential to be broadly neutralizing, deemed bnAbs.^14,35,36^ We compared the H2 vaccine-elicited RBS mAbs with known neutralizing RBS mAbs to identify and dissect molecular features associated with broadly neutralizing activity. Previously, we observed the reliance of mAb 1-1-1F05’s interaction with HA on its long 20 amino acid CDRH3 loop (Fig. 4C and 6D, left). This interaction resembles that of bnAb C05, and structural alignment of the two reveals a near identical topology at the interface with both CDRH3 loops interacting with the RBS through a hydrogen bonding network (Fig. 6D, left). Another common feature of H2-specific neutralizing RBS mAbs is the presence of an aromatic residue that interacts with the conserved W153 of HA. This aromatic motif is found regularly in VH1-69 encoded mAbs—often as F54 in the CDRH2 loop or as another aromatic residue within the CDRH3 loop—and forms a crucial interaction with the tryptophan residing in the RBS’s hydrophobic cavity.^36^ MAb 1-1-1E04, which is also encoded by the VH1-69 gene, interacts in a similar manner with F112b of its CDRH3 loop. Structural alignment of 1-1-1E04 with other VH1-69-encoded mAbs 2G1 (PDB: 4HG4), 8M2 (PDB: 4HFU), and 8F8 (PDB: 4HF5) demonstrate the similarity in aromatic interactions with HA’s W153 (Fig. 6D, middle). Notably, the VH4-59-encoded 1-1-1F05 lacks this signature aromatic interaction.

In contrast to the cross-reactive RBS antibodies, the H2-specific mAb 4-1-1E02, encoded by the VH 4-61 gene, uses a large footprint with substantial contributions from its heavy and light chains (Fig. 6C and D, right). Crucial interactions included P100 on its CDRH3 loop binding in the hydrophobic pocket, favorable electrostatic interactions between N33 in the CDRH1 loop and E190 of HA, and presumed hydrogen bonding between Y92 in the CDRL3 loop with N144 of HA.

Overall, H2 vaccination elicited novel antibodies that targeted a variety of epitopes including the RBS and antigenic site Sa. Moreover, multiple cross-reactive mAbs targeted the RBS with similar mechanisms as known RBS-targeting mAbs, suggesting H2 vaccination offers a broad scope of protection by generating multiple modes of binding to the RBS.

### Novel ‘medial junction’ epitope targeted by H2 vaccine-elicited antibodies

In addition to RBS-targeting mAbs, we also observed cross-reactive antibody responses to non-RBS epitopes. Particularly, the medial junction epitope was targeted, which encompasses the region between the conserved central helix and vestigial esterase domain of HA (Fig. 4A and B). As a polyclonal response to the medial junction epitope was not observed by EMPEM at week 0, we expect the pAbs binding at this epitope were boosted by H2 vaccination.

We investigated this novel epitope further by isolating cross-reactive mAb 2-2-1G06, which binds at the medial junction of H1, H2, and H5 strains (Fig. 4C). While this epitope has not been previously shown to be targeted in influenza A viruses, it resembles influenza B virus bnAb CR8071.^37^ To investigate the molecular interactions important for 2-2-1G06’s broad reactivity, we obtained cryoEM structures of 2-2-1G06 in complex with H2 HA (2.9Å resolution; Fig. 7A) and H1 HA (3.1Å resolution; Fig. S8A). MAb 2-2-1G06 utilizes a broad footprint with interactions from all its CDR loops to mediate binding (Fig. 7B and Fig. S8B–E). The strongest interactions were within the CDRH3 and CDRL2 loops. Residue Y100A, which resides at the top of the CDRH3 loop, inserts into the interface area and forms a cation-ρε interaction with D419 of the HA’s highly conserved central helix. At the turn of framework region 3, R68 faces the upper central helix of HA and interacts electrostatically with HA’s E407 (Fig. 7C).

**Fig. 7.**
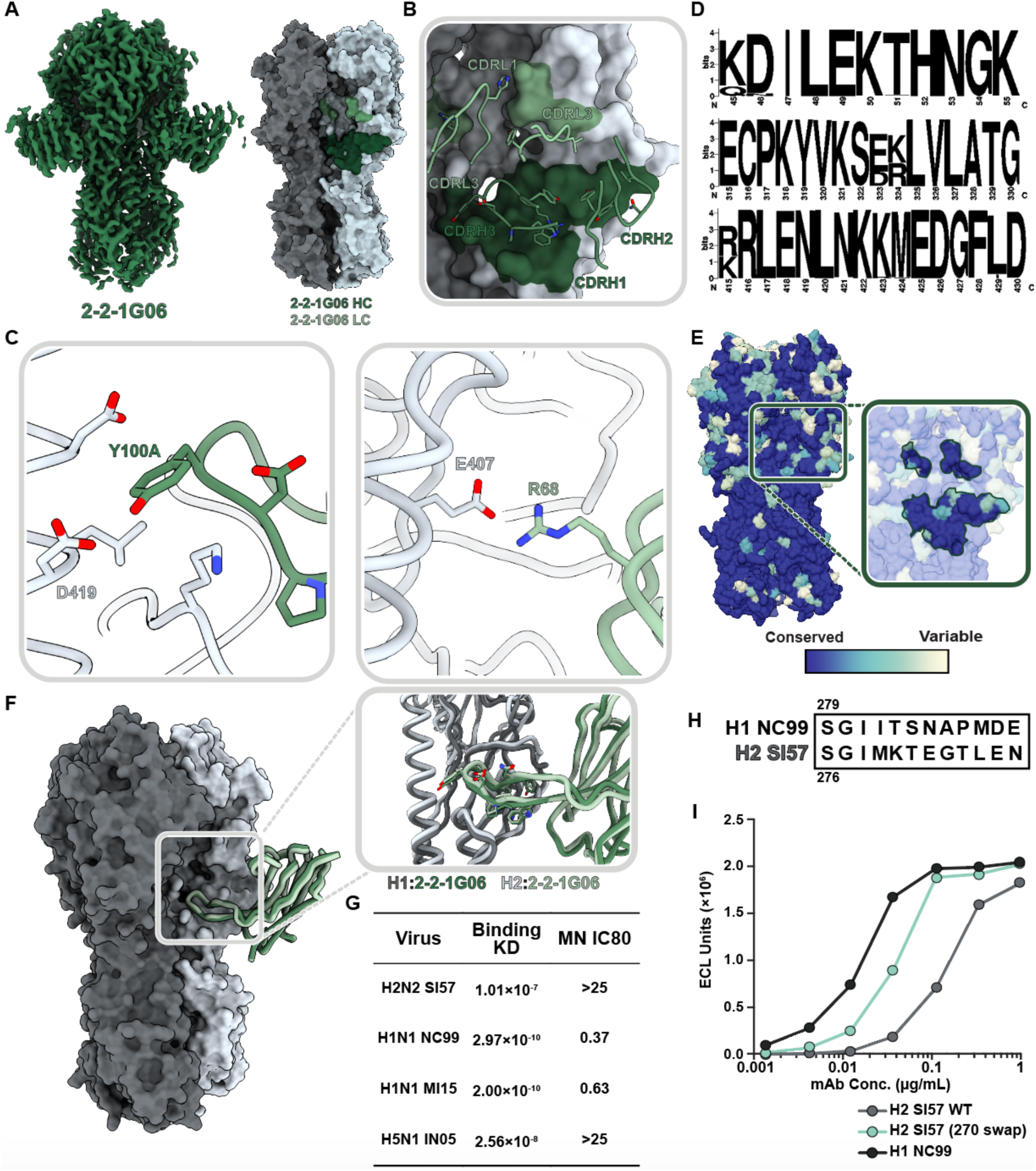
Structural and functional characterization of 2-2-1G06 targeting the novel ‘medial junction’ epitope. (A) CryoEM map of 2-2-1G06 in complex with H2 HA (left) and antibody footprint (right). (B) CDR loop interactions at the 2-2-1G06 epitope. (C) 2-2-1G06 epitope-paratope interactions. Residues presumed critical for binding are shown (Y106 of the CDRH3 on the left and R68 of the CDRL2 on the right). (D) Sequence alignment of 180 human and avian H2 viruses. (E) 16 years of H1 HA sequence variability mapped on an HA surface. Years with sequences represented include 1999, 2006, 2007, 2008, 2009, 2011, 2013, and 2015. (F) Structural comparison of 2-2-1G06 in complex with H2 and H1 NC99. Pop-out panel shows CDRH3 residues. (G) 2-2-1G06 binding affinity and microneutralization of H1, H2, and H5 virus. (H) 270 loop sequence alignment of H1 and H2 strains used in neutralization assay. (I) Binding activity of 2-2-1G06 to SI57 H2 WT, H2 with H1-reverted mutations “270 swap,” and H1 NC99.

The medial junction epitope is highly conserved across Group 1 subtypes as evidenced by its sequence homology among 180 sequence-aligned H2 strains (Fig. 7D) and its near universal conservation within the past 16 years for H1 viruses (Fig. 7E) The epitope-paratope interface and binding topology between 2-2-1G06 and HA is structurally near-identical between H1 and H2 subtypes (Fig. 7F). Despite 2-2-1G06’s ability to neutralize pre- and post-pandemic strains of H1 virus and cross-reactivity to H1, H2, and H5 HA, it was unable to neutralize H2 and H5 virus strains. We expect this may relate to 2-2-1G06’s slower on-rate and pronounced off-rate to H2 and H5 compared to H1 (Fig. 7G; Fig. S8F).

To dissect the molecular distinctions between H1 and H2 (Fig. 7H) that lead to differences in 2-2-1G06 binding and neutralization, we assessed the contribution of H1’s 270 loop residues, which are positioned near the apical edge of the 2-2-1G06 epitope (HA residues 265-276). We generated an H2 HA mutant with positions 265-276 mutated to H1 residues (Fig. 7H; strain A/New Caledonia/20/1999; NC99). This mutant, H2 (270 swap), saw restored binding of 2-2-1G06, though it did not reach the potential of WT H1 (Fig. 7I). Based on these data, we suspect the 270 loop residues provide ancillary—yet not complete—support to the overall binding potential of mAb 2-2-1G06 to H2.

Overall, these studies identify an unappreciated neutralizing epitope on the medial junction of HA that is targeted by pAbs post H2 vaccination and shows potential for broad reactivity and neutralization.

## DISCUSSION

Human immune responses to seasonal influenza virus infection tend to be biased toward variable, strain-specific epitopes on the HA head, providing motivation to create vaccines that redirect B-cell responses to conserved sites on HA. Understanding the interplay and context of antibody responses is crucial for developing vaccine regimens that boost desirable responses and limit strain-specific recall in people of different age groups. Recently, vaccinations with an experimental H2-F and H2 DNA plasmid vaccines were shown to induce H2-specific antibodies as well as bnAbs targeting the central stem epitope in H2-naïve human populations.^27,28^ Human clinical trial data now provide an opportunity to further refine this promising approach and advance it toward a more universal vaccine. Here, we investigated the proportion and dynamics of pAb responses to the H2 vaccines.

Our work explores the complex immune dynamics of pre-exposed and naïve individuals post-vaccination. Primary exposure to H2N2 viruses through vaccination with H2 DNA plasmid or H2-F vaccine candidates elicited pAb responses to conserved epitopes on HA. For H2-naïve individuals, we observed through semi-quantitative nsEMPEM and serological analyses that the vast majority of H2-specific pAb responses targeted the conserved stem domain, and H1 cross-reactive RBS responses were also recalled. Moreover, secondary exposure through H2 vaccination in pre-exposed individuals or H2-F boost in naïve individuals expanded the diversity of pAb responses to target variable head epitopes, shifting the dominance of pAb landscapes to the head domain. These results demonstrate the dynamics of first recalling cross-reactive memory B cells upon exposure to novel influenza virus strains while eliciting more strain-specific naïve B cells, a process that amplifies upon re-exposure to the same virus.

We also note stark differences between participant outcomes based on the first vaccine dose employed. Through our semi-quantitative EMPEM analyses, we show the proportion of H2 HA molecules bound to pAbs increased by over 70% post-DNA plasmid primary vaccine dose (week 4) to post H2-F boost in participant 1-1 (week 20). In general, naïve participants who received an H2-F first dose showed quicker increases in pAb-bound HA (from week 0 to week 4) with less dramatic increases after H2-F boost. Regardless of first vaccine dose, H2-naïve participants showed antibody responses that diversified from being almost wholly stem-targeting post-first-dose to being dominated by head responses after H2-F boost. Taken together, these data demonstrate the importance of primary vaccine dose on pAb outcomes and the polarizing phenotype of primary and secondary exposures to HA.

While the majority of universal vaccine efforts focus on the HA stem domain, conserved epitopes in the HA head domain remain promising for their ability to potently neutralize receptor binding and inhibit viral entry. The DNA plasmid and H2-F vaccines induced cross-reactive, neutralizing antibodies to the RBS in multiple participants that circulated as pAbs in serum. Moreover, H2 vaccination induced cross-reactive antibodies with diverse gene usages and mechanisms of binding, providing redundancy that may safeguard against viral escape mutations. In contrast, a strain-specific mAb elicited by secondary exposure to the vaccine had a larger footprint that extended into antigenic site Sa. These results suggest pinpointing minimal conserved HA epitopes and key residues is crucial for expanding breadth of Ab responses.

Novel, conserved epitopes on HA are still being discovered, such as the anchor, trimer interface, and lateral patch epitopes,^9,29,38–43^ suggesting that there are more avenues to exploit for universal vaccine design. Here, we describe the novel influenza A medial junction epitope targeted by pAbs and describe its molecular details with cross-reactive mAb 2-2-1G06. This epitope resides in the cavity of the stem’s central helix and head domain junction. Though this marks the first description in influenza A viruses, Abs to a similar epitope inhibit release of progeny virions of influenza B viruses. In an H2-naïve participant, we observed recall of H1-reactive medial junction cavity pAbs upon first exposure to H2. While we did not observe medial junction pAbs against H2 in serum via EMPEM, mAb 2-2-1G06 isolated from a memory B cell population of the same donor bound H1, H2, and H5 *in vitro*. Additionally, it was able to neutralize pre- and post-2009 H1N1 pandemic strains. These results suggest H2 vaccination can induce memory B cells generated by H1N1 exposure that recall antibodies to the cross-reactive medial junction cavity in H2.

Due to the ability of influenza virus to subvert antibody responses, targeting protective epitopes and limiting person- and context-dependent variation is crucial for developing vaccines targeting diverse influenza strains regardless of exposure status. This vaccine trial provides strong proof-of-concept that H2 vaccination can induce cross-reactive pAbs to conserved epitopes in the RBS and central stem; however, we demonstrate that targeting conserved epitopes is substantially reduced if there is pre-existing immunity to H2. Our results will help inform modifications to H2 HA immunogens and prime-boost strategies to further improve vaccine efficacy and breadth regardless of exposure history.

## ACKNOWLEDGEMENTS.

The authors thank Bill Anderson and Hannah Turner from the Scripps Research Institute for their help with electron microscopy experiments. We thank Dr. Lauren Holden for her help in preparing this manuscript. This work is supported by the Bill and Melinda Gates Foundation through grant INV-004923 (to ABW). JH is funded by NIAID 2 T32 AI007244-36. HRP is supported by a David C. Fairchild Endowed Fellowship, the Achievement Rewards for College Scientists Foundation, and NIH F31 Ruth L. Kirschstein Predoctoral Award 1F31Al172358. RAG, SAC, JER, LYC, AC, MK, KVH, GLC, SFA and ABM are supported by funding from the National Institutes of Health (NIH) Intramural Research Program.

## AUTHOR CONTRIBUTIONS

Conceptualization: JH, YRY, SFA, ABW, ABM

Methodology: JH, YRY, SFA

Formal analysis: YRY, JH, SFA, HRP, STR, AMJ, AJR, RAG, SAC, JER, LYC, AC

Investigation: YRY, JH, SFA, STR, AMJ, AJR

Resources: ABW, ABM, GLC, KVH

Writing of original draft: JH, YRY, SFA, HRP

Reviewing, editing, and other feedback: ABW, ABM

Visualization: HRP, YRY, JH, SFA

Supervision: ABW, ABM, MK

Project admin: JH, SFA

Funding acquisition: ABW, ABM

## DATA AVAILABILITY

Maps generated from the electron microscopy data are deposited in the Electron Microscopy Databank (http://www.emdatabank.org/) under accession IDs EMD-41464, EMD-41465 EMD-41466 EMD-41467 EMD-41468 EMD-41469 EMD-41470 EMD-41471 EMD-41472 EMD-41473 EMD-41474, EMD-41514, EMD-41515, EMD-41516, EMD-41517, EMD-41518, EMD-41519, EMD-41520, EMD-41521, EMD-41522, EMD-41523, EMD-41524, EMD-41525, EMD-41526, EMD-41527, EMD-41528, EMD-41529, EMD-41530, EMD-41531, EMD-41532, EMD-41533, EMD-41534, EMD-41535, EMD-41536, EMD-41537, EMD-41538, EMD-41539, EMD-41540, EMD-41541, EMD-41542, EMD-41543, EMD-41544, EMD-41545, EMD-41546, EMD-41547, EMD-41548, EMD-41549, EMD-41550, EMD-41551, EMD-41552, EMD-41553, EMD-41554, EMD-41555, EMD-41556, EMD-41557, EMD-41558, EMD-41559, EMD-41560, EMD-41561, EMD-41562, EMD-41563, EMD-41564, EMD-41683, EMD-41684, EMD-41685, EMD-41686, EMD-41687, EMD-41688, EMD-41689, EMD-41690, EMD-41691, EMD-41692, EMD-41693, and EMD-41694. See Tables S1, S2, and S3 for more details. Atomic models corresponding to these maps have been deposited in the Protein Data Bank (http://www.rcsb.org/) under accession IDs 8TP2, 8TP3, 8TP4, 8TP5, 8TP6, 8TP7, 8TP9, and 8TPA.

## METHODS

### Selection of clinical participants

The VRC 316 clinical trial was a phase I, open-label, and randomized (ClinicalTrials.gov, NCT03186781) and has been described previously.^27^ In short, the study was conducted at the National Institutes of Health (NIH) Clinical Center by the Vaccine Research Center Clinical Trials Program of the National Institute of Allergy and Infectious Diseases (NIAID). Trial protocols were approved by the NIAID institutional review board and informed consent was obtained from each enrolled participant. The trial evaluated H2 vaccination with H2 plasmid DNA encoding H2 A/Singapore/1/1957 or homologous H2 HA Ferritin nanoparticle followed by a boost 16 weeks later in all participants with the H2 HA Ferritin nanoparticle vaccine. For each of the two vaccine regimens, participants born before 1966 (H2 pre-exposed) or after 1969 (H2 naïve) were enrolled for a total of 4 vaccine groups. Twelve representative participants, 3 from each of the trial groups were chosen for ad-hoc analysis in this study.

### HA expression and purification

HA proteins were transiently expressed in HEK 293F cells (Thermo Fisher) at a density of 1.0 x 10^6^ cells/mL with a 1:3 ratio of DNA to PEIMax. HEK 293F cells were maintained in 293FreeStyle expression medium (Life Technologies) and cultured at 37°C, 8% CO_2_, and shaken at 125 rpm. Six days after transfection, cells were harvested and spun down. HAs were purified by a HisTrap column (Cytiva). After elution, HA trimers were purified by size exclusion chromatography using a Superdex 200 Increase 10/300 column (GE Healthcare). Fractions corresponding to trimeric HA were pooled, concentrated, and buffer exchanged to TBS using 50 kDa Amicon concentrators.

### Polyclonal antibody purification and digestion to pFabs

Serum samples were collected from participants in the H2 vaccine clinical trial (NCT03186781).^27^ Serum samples were first heat inactivated in a 55°C water bath for 30 minutes. Inactivated serum was incubated for 20 hrs with protein G (GE Healthcare) or CaptureSelect resin (Thermo Fisher) at a ratio of 1 mL serum to 1 mL resin slurry. Samples were centrifuged briefly and IgG-depleted serum removed. The IgG-rich resin was washed three times with PBS by centrifugation. IgG was eluted from resin after incubating with 0.1 M glycine, pH 2.5 buffer for 20 minutes. The eluent was then neutralized with a 1 M Tris-HCl pH 8.0 buffer. The solution was buffer exchanged to PBS using 50 kDa Amicon concentrators.

Next, purified IgGs were digested to pFabs. Papain (Sigma Aldrich) was activated in fresh digestion buffer (20 mM sodium phosphate, 10 mM EDTA, 20 mM cysteine at pH 7.4). We incubated 4 mg of polyclonal IgG with activated, immobilized papain for 18-22 hours at 37°C. Digested IgG was separated from papain using Pierce spin columns (Thermo Fisher) and buffer exchanged to TBS using 50 kDa Amicon concentrators. We separated pFab and Fc from undigested IgG using size exclusion chromatography with a Superdex 200 Increase 10/300 column (GE Healthcare) and concentrated the pFab/Fc.

### Monoclonal antibody digestion and Fab purification

Fabs of monoclonal antibodies were generated by papain digestion of purified IgG. Papaya latex papain (Sigma Aldrich) was activated in a fresh solution of 20 mM sodium phosphate, 10 mM EDTA, and 20 mM cysteine at pH 7.4 for 15 minutes at 37°C. IgG was digested in the activated papain solution for 4 hours at 37°C in a ratio of 1 mg IgG to 40 μg papain. The reaction was quenched using 50 mM iodoacetamide. The digestion products were buffer exchanged to PBS via centrifugation with 30 kDa Amicon concentrators, purified on a Superdex 200 Increase 10/300 column (GE Healthcare), and concentrated using 30 kDa Amicon concentrators.

### HA complex formation with pFabs and monoclonal Fabs

PFab-HA complexes were obtained by incubating 500 μg concentrated pFab/Fc mixture with 10 μg recombinant HA at room temperature for 16-20 hours. Complexes were purified from unbound HA, pFab, and Fc using size exclusion chromatragraphy with a Superdex 200 Increase 10/300 column (GE Healthcare) and concentrated using 100 kDa Amicon concentrators. Monoclonal Fabs were incubated with HA at a 3:1 molar ratio for 1 hour at room temperature.

### Negative stain electron microscopy

Immune complexes were deposited on glow-discharged (PELCO easiGlow, Ted Pella, Inc.) carbon-coated 400 mesh copper grids (Electron Microscopy Sciences) at a concentration of approximately 20 μg/mL. Excess sample was removed by blotting, and grids were stained with two back-to-back depositions of 2% w/v uranyl formate for 60 s each. Excess stain was removed by blotting and the grids were allowed to dry.

Grids were imaged on either a 200 kV Tecnai F20 electron microscope (FEI) with a TemCam F416 CMOS camera (TVIPS) or a 120 kV Tecnai Spirit T12 (FEI) with an Eagle CCD 4k camera (FEI). Images were collected at 62,000 or 52,000X magnification with pixel sizes of 1.77 and 2.06 Å, respectively. Micrographs were acquired using the Leginon software package and Appion was used to pick 100,000-400,000 single particles.^44–46^ Particles were then processed to reference-free 2D class averages and 3D reconstructions using Relion.^47–49^ UCSF Chimera and UCSF ChimeraX were used to analyze data and generate figures.^50,51^ Due to limitations such as low particle count for rare polyclonal specificities and lack of angular sampling, some epitope specificities were not amenable to 3D reconstruction but showed clear specificity to HA epitopes as confirmed by distinct 2D class averages, as has been observed previously.^52^ When applicable, these specificities are shown as flat surface colors with dotted black outlines.

### Semi-quantitative analysis of nsEMPEM data

After the first round of reference-free 2D class averaging of polyclonal samples post-EM imaging, all classes with HA were selected, removing junk particles. A second round of reference-free 2D class averaging was performed on the selected particle stack. For semi-quantitative analysis, only side view particles were counted, as it is not possible to distinguish epitope specificities with top views. Classes were grouped according to number of pFabs bound to HA (0-4 pFabs/HA) and particle counts from Relion^48^ were recorded for each group. As 2D classification is often inadequate to assign overlapping and neighboring epitopes, pFab specificities to the head and stem domains were also grouped and counted, rather than individual specificities. Particle counts for pFab abundance and head/stem specificity were conducted on the same particle stacks independently.

### MSD binding assay

Meso Scale Discovery (MSD) 384 well Streptavidin coated SECTOR Imager 6000 Reader Plates were blocked with 5% MSD Blocker A for 30 to 60 minutes, then washed six times with the wash buffer (PBS+0.05% Tween). The plates were then coated with biotinylated HA protein (same protein as was used for flow cytometry) for one hour and washed. mAbs were diluted in 1% MSD Blocker A to 1μg/ml, serially diluted three-fold, and added to the coated plates. Serum samples were diluted 1:100 in 1% MSD Blocker A and serially diluted three-fold before adding to coated plates. A control mAb (53-1F12)(Andrews et al., 2017) was added to each plate to use a reference standard for each assay. After a one hour incubation with sera or mAbs, plates were washed and incubated for one hour with SULFO-TAG conjugated anti-human IgG for mAbs (MSD) or SULFO-TAG conjugated polyclonal anti-human IgG+A+M (Thermo Fisher) for serum samples. After washing, the plates were read using 1X MSD Read Buffer using a MSD SECTOR Imager 600. For mAbs, binding curves were plotted and the area under the curve (AUC) was determined using GraphPad Prism 8. For sera, binding of 1 μg/mL of 53-1F12 to H2 or H2 stabilized stem was assigned a concentration of 100 arbitrary units per milliliter (AU/mL). Serial dilutions of sample within the dynamic range of the standard curve were interpolated to assign a sample concentration in AU/mL. Results were plotted and analyzed using GraphPad Prism 8.

The following HA strains were used for mAb and/or serum binding assays: H1 A/NewCaledonia/20/1999 (NC99) ectodomain, H1 A/Michigan/45/2015 (MI15) ectodomain, H5 A/Indonesia/05/2005 (IN05) ectodomain, H2 A/Singapore/2/1957 (SI57) ectodomain, H2 SI57 ectodomain with 270 swap, H2 SI57 stabilized stem, H6 A/Taiwan/1/2013 (TW13) ectodomain, and H9 A/Hongkong/1073/1999 (HK99) ectodomain, and H3 A/Hongkong/4801/2014 (H3 HK14).

### Single-cell sorting HA-specific B cells

Cryopreserved PBMCs from blood collected 1 and 2 weeks after the H2-F boost were stained with anti-human monoclonal antibodies CD3 BV510 (OKT3, 1:400 dilution, BioLegend, RRID:AB_2561376), CD56 BV510 (HCD56, 1:200 dilution, BioLegend, RRID:AB_2561385), CD14 BV510 (M5E2, 1:200 dilution, BioLegend, RRID:AB_2561379), CD27 BV605 (O323, 1:50, BioLegend, RRID:AB_11204431), CD20 APC-Cy7 (2H7, 1:400 dilution, BioLegend, RRID:AB_314261), IgG BV421 (G18-145, 1:50 dilution, BD Biosciences, RRID:AB_2737665), IgM PercpCy55 (G20-127, 1:40 dilution, BD Biosciences, RRID:AB_10611998), CD19 ECD (H3-119, 1:50 dilution, BD Biosciences, RRID:AB_130854), CD21 PeCy5 (B-ly4, 1:100 dilution, BD Biosciences, RRID:AB_394028) and CD38 (HIT2, 1:400 dilution, BD Biosciences, RRID:AB_1727472). H2 A/Singapore/2/1957 ectodomain and stabilized stem HA probes were expressed, biotinylated and labeled with fluorochromes as described previously (Whittle et al., 2014). Aqua dead cell stain was added for live/dead discrimination (ThermoFisher Scientific). Stained samples were run on a FACS Aria II (BD Biosciences) and data analyzed using FlowJo (TreeStar). CD3- CD14- CD56- CD19+ CD20- CD21-CD27^hi^ CD38^hi^ plasmablasts or CD3- CD14- CD56- CD19+ CD20+ IgG+ IgM-Memory B cells were gated, and H2 HA-binding B cells were single-cell sorted into 96-well plates. H2 HA head-specific B cells were identified by indexing.

### Single-cell Ig amplification and sequencing and mAb production

Reverse transcription was performed on sorted cells and multiplexed PCR was used to amplify immunoglobulin heavy and light chain genes as described previously.^53,54^ We obtained paired heavy and light chain Ig sequences from an average of 70% of single cells on which we performed PCR. PCR products were sequenced by Beckman Coulter or Genewiz.

Heavy and light chain sequences were synthesized and cloned by Genscript into IgG1, kappa, or lambda expression vectors. To produce mAbs recombinantly, Expi293 cells were transfected with plasmids encoding Ig heavy and light chain pairs with ExpiFectamine (ThermoFisher Scientific). Monoclonal antibodies were purified from the cell supernatant using sepharose Protein A (Pierce).

### CryoEM grid preparation and imaging

Immune complexes were prepared as described above and applied to grids at a concentration of 0.4-0.8 mg/mL. Octyl-beta-glucoside detergent was added to samples at a final concentration of 0.1% immediately before deposition on glow-discharged Au 1.2/1.3 400-mesh and 2/2 200 mesh grids (Electron Microscopy Services). Samples were incubated on grids for 7 seconds before being blotted off and plunge-frozen in liquid ethane using a Vitrobot mark IV (Thermo Fisher).

After freezing, cryo grids were loaded into a 300 kV FEI Titan Krios or 200 kV Talos Arctica (Thermo Fisher), both of which were equipped with K2 Summit direct electron detector cameras (Gatan). Data were collected with approximate exposures of 50 e^-^/Å^2^. Magnifications of 130,000 or 36,000X were used for the Krios or Arctica, respectively. Data collection was automated using Leginon. Further details are described in Table S1.

### CryoEM data processing

Image pre-processing was performed with the Appion software package.^45^ Micrograph movie frames were aligned, dose-weighted using the UCSF MotionCor2 software,^55^ and GCTF was estimated.^56^ Micrographs were then transferred to CryoSPARC v3.0 for particle picking and reference-free 2D classification.^57^ Initial 2D classes of high quality were used as templates for template picking of datasets followed by 2D classifications to remove bad particles. Global 3D refinements were performed, and particle stacks were sorted for Fab-bound complexes by heterogeneous refinements and 3D variability analyses.^58^ Some datasets were further analyzed in Relion, where they were sorted using alignment-free 3D classification.

For polyclonal samples, 40 Å sphere masks were used to separate particles with pAbs bound within the masked area by 3D Variability (CryoSPARC) or through alignment-free 3D classification (Relion). Once pAb complexes were separated, they were refined separately and new masks featuring the full immune complex were used for final refinements.

More details are described in Table S1 including imposed symmetry and final particle counts. Figures were made using UCSF Chimera and ChimeraX.

### Atomic model building and refinement

For monoclonal EM maps, we refined atomic models using corresponding post-processed maps. PDBs 6CF7 and 2WR7 were used as the initial H1 and H2 models, respectively. PDB 6CF7 was mutated to the A/New Caledonia/20/1999 sequence. Initial models for Fabs were predicted using ABodyBuilder.^59^ Both HA and Fab models were manually fit into density using Coot.^60^ Iterative manual model building in Coot followed by Rosetta^61^ relaxed refinements were used to generate atomic models of each complex. We evaluated our models using MolProbity and EMRinger of the Phenix software package^62–64^ and the PDB validation server. Epitope-paratope interactions were analyzed and visualized in UCSF Chimera and ChimeraX. Models are numbered based on the H3 numbering system for HA and the Kabat numbering system for Fabs.

### Sequence alignment and conservation assessment

H1 HA sequence variability was assessed based on 8 distinct human H1 strains from 1999, 2006, 2007 2008, 2009, 2011, 2013, and 2015. The conservation model was generated using sequence logo in the Librator^65^ application and visualized in UCSF Chimera. A survey of 180 H2 HA sequences from human and avian viruses was conducted using sequences from the Influenza Research Database^66^ and represented as a sequence logo using the web logo tool.^67^

### Microneutralization assay

Generation of the replication-restricted reporter (R3ΔPB1) virus H1N1 and Rewired R3ΔPB1 (R4ΔPB1) virus H2N2 has been described elsewhere.^68^ Briefly, to generate the R3/R4ΔPB1 viruses the viral genomic RNA encoding functional PB1 was replaced with a gene encoding the fluorescent protein (TdKatushka2), and the R3/R4ΔPB1 viruses were rescued by reverse genetics and propagated in the complementary cell line which expresses PB1 constitutively. Each R3/R4ΔPB1 virus stock was titrated by determining the fluorescent units per mL (FU/mL) prior to use in the experiments. For virus titration, serial dilutions of virus stock in OptiMEM + TPCK were mixed with pre-washed MDCK-SIAT-PB1 cells (8 x 105 cells/ml) and incubated in a 384-well plate in quadruplicate (25 µL/well). Plates were incubated for 18-26 h at 37°C with 5% CO2 humidified atmosphere. After incubation, fluorescent cells were imaged and counted by using a Celigo Image Cytometer (Nexcelom) with a customized red filter for detecting TdKatushka2 fluorescence.

For the microneutralization assay, serial dilutions of antibody were prepared in OptiMEM and mixed with an equal volume of R3/R4ΔPB1 virus (∼8 x 104 FU/mL) in OptiMEM + TPCK. After incubation at 37°C and 5% CO2 humidified atmosphere for 1 h, pre-washed MDCK-SIAT-PB1 cells (8 x 105 cells/well) were added to the serum-virus mixtures and transferred to 384-well plates in quadruplicate (25 µL/well). Plates were incubated and counted as described above. Target virus control range for this assay is 500 to 2,000 FU per well, and cell-only control is acceptable up to 30 FU per well. The percent neutralization was calculated for each well by constraining the virus control (virus plus cells) as 0% neutralization and the cell-only control (no virus) as 100% neutralization. A 7-point neutralization curve was plotted against serum dilution for each sample, and a four-parameter nonlinear fit was generated using Prism (GraphPad) to calculate the 80% (IC80) inhibitory concentrations.

### Biolayer interferometry

Biolayer interferometry was performed with an Octet Red384 (FortéBio). Biotinylated HA protein (A/New Caledonia/20/99, A/Michigan/45/2015, A/Indonesia/05/2005, A/Singapore/1/1957) at 5 µg/mL in assay buffer (PBS + 1% BSA), was loaded onto a PBS buffer equilibrated streptavidin-coated biosensor (Sartorius) for 300 seconds. Biosensors were then equilibrated with assay buffer to remove unbound HA protein to establish a baseline signal for 15 seconds. Once the baseline was determined, HA protein bound biosensors were assigned to different concentrations of Fab (1600 nM, 400 nM, 100 nM, and 25 nM). After 180 seconds of association, HA-Fab complexed biosensors were transferred to baseline wells, measuring dissociation for 300 seconds.

