## Supplemental Information for "Immune memory shapes human polyclonal antibody responses to H2N2 vaccination"

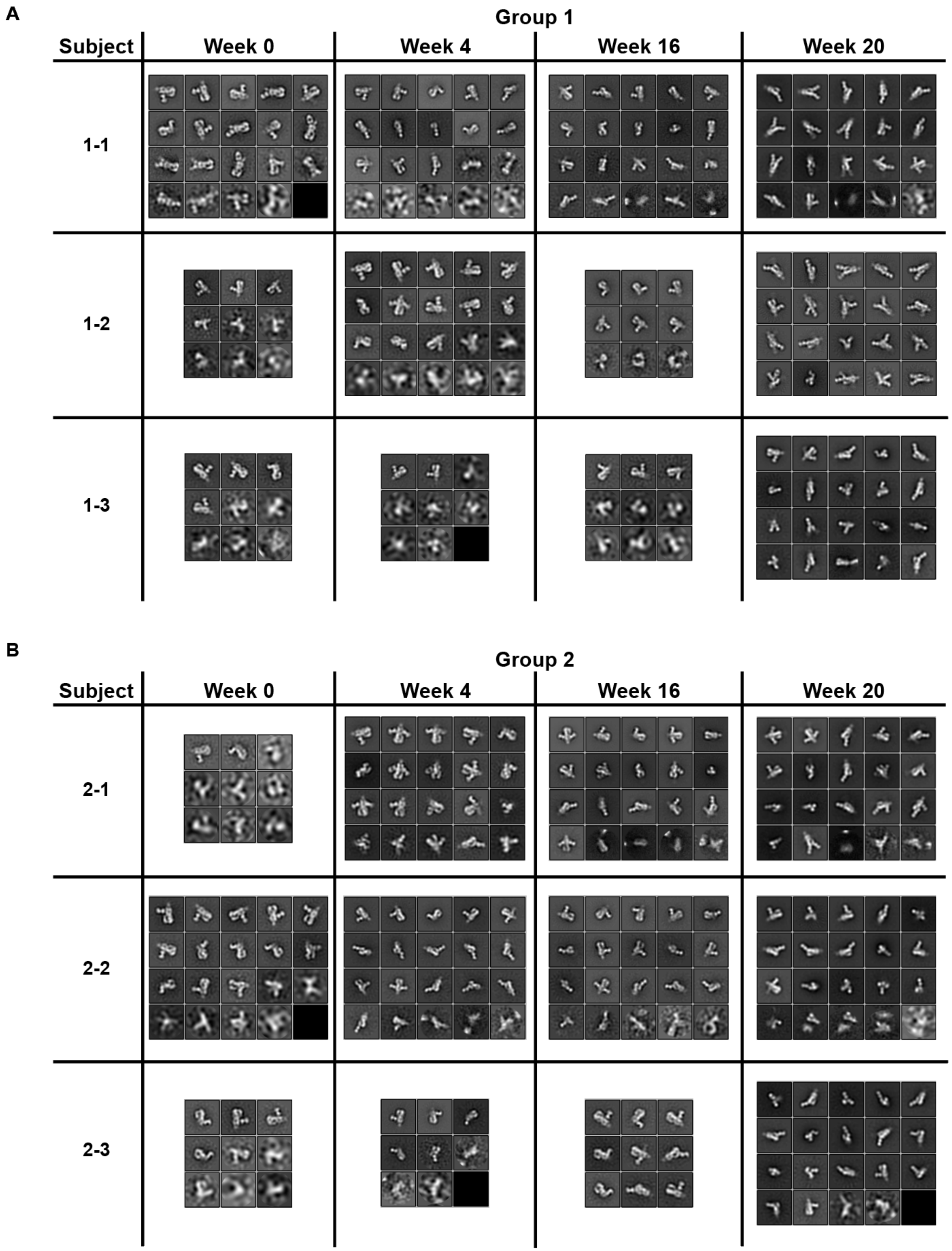


**Figure S1: Group 1 and 2 sample 2D classes**. Related to Figure 2. Sample 2D classes that make up 3D models shown in Figure 2 for Group 1 (A) and Group 2 (B). Datasets with >7.3k particles are divided into 20 classes while those with <7.3k particles are divided into 9. All 2D classification datasets are shown in order of descending particle count.

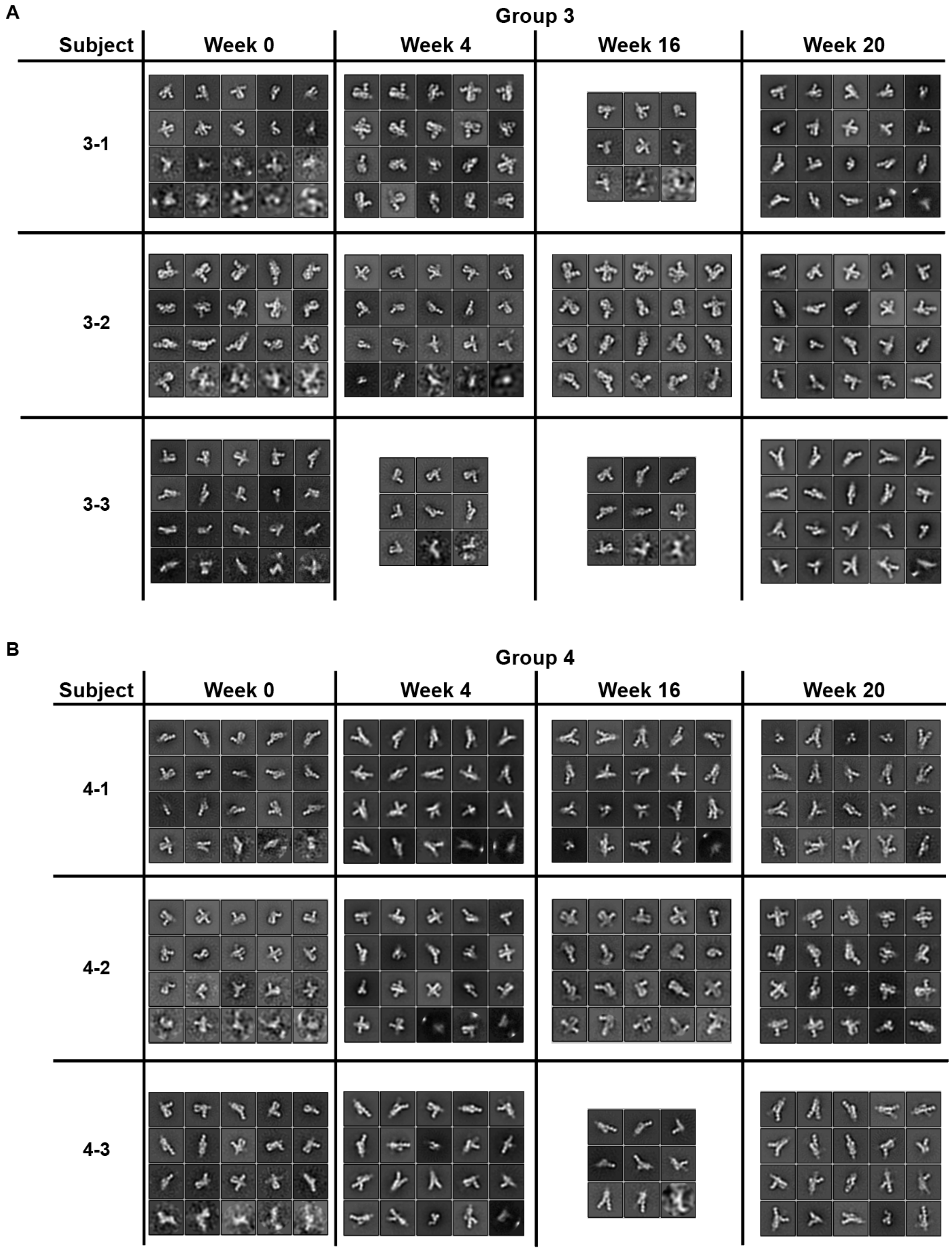


**Figure S2: Group 3 and 4 sample 2D classes**. Related to Figure 2. Sample 2D classes that make up 3D models shown in Figure 2 for Group 3 (A) and Group 4 (B). Datasets with >7.3k particles are divided into 20 classes while those with <7.3k particles are divided into 9. All 2D classification datasets are shown in order of descending particle count.

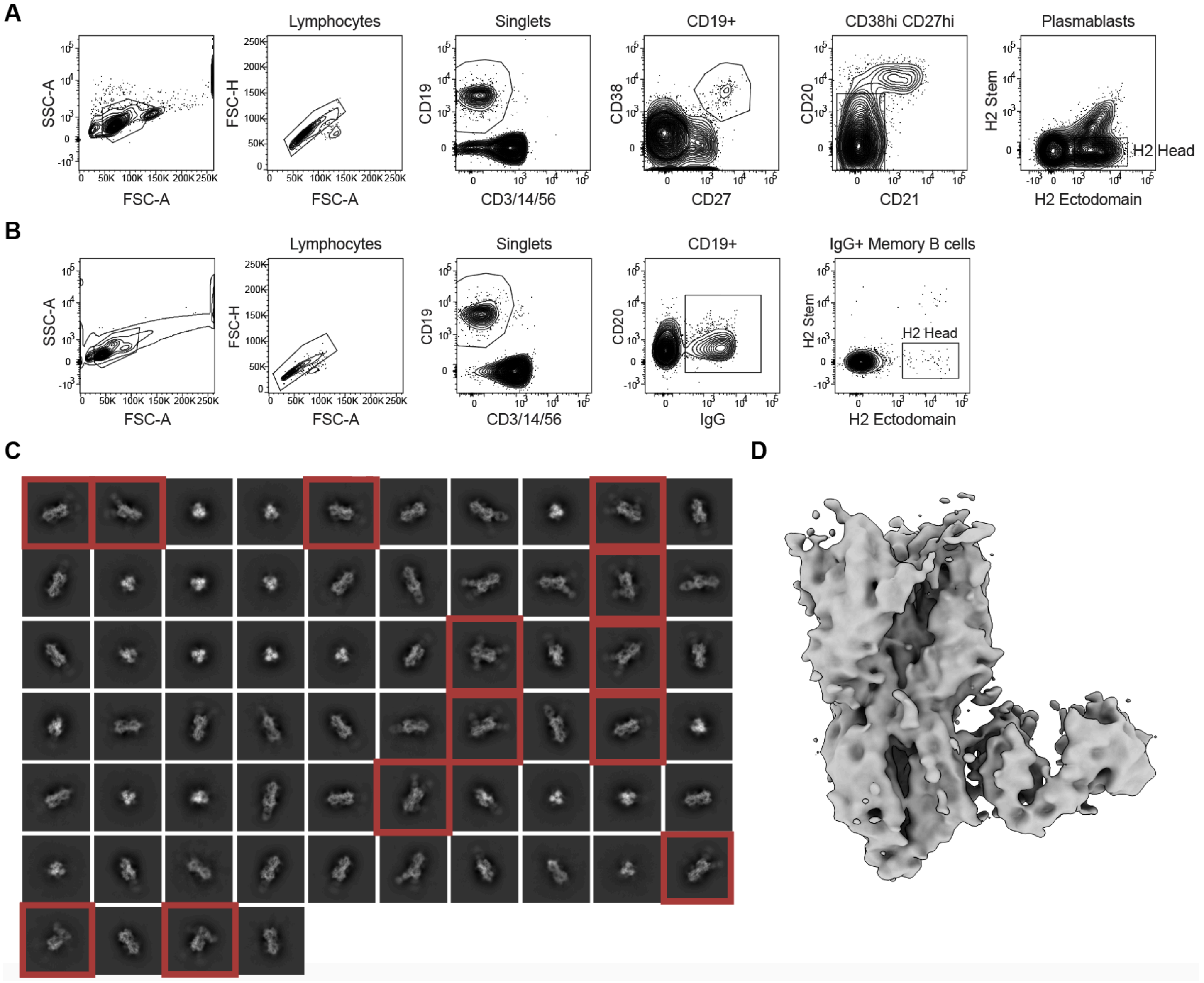


**Figure S3: B-cell sorting strategy and polyclonal stem responses observed by cryo-EM**. Related to Figure 4 and 5. Flow cytometry gates used to detect and sort H2 HA head-specific B cells from the CD19+ CD3/14/56 (dump)- CD27$^{hi}$ CD38$^{hi}$ CD20$^{lo}$ CD21$^{lo}$ plasmablast (A) or CD19+ CD3/14/56 (dump)- CD20+ IgG+ memory B cell compartment (B). Each plot shows the cell population gated immediately to the left as indicated above each plot. H2 HA head-specific B cells were detected as H2 HA ectodomain+ H2 stem-. (C) 2D classes of HA bound to polyclonal antibodies. Classes featuring pFabs with stem specificities outlined in red. (D) 3D reconstruction of stem-specific pFab.

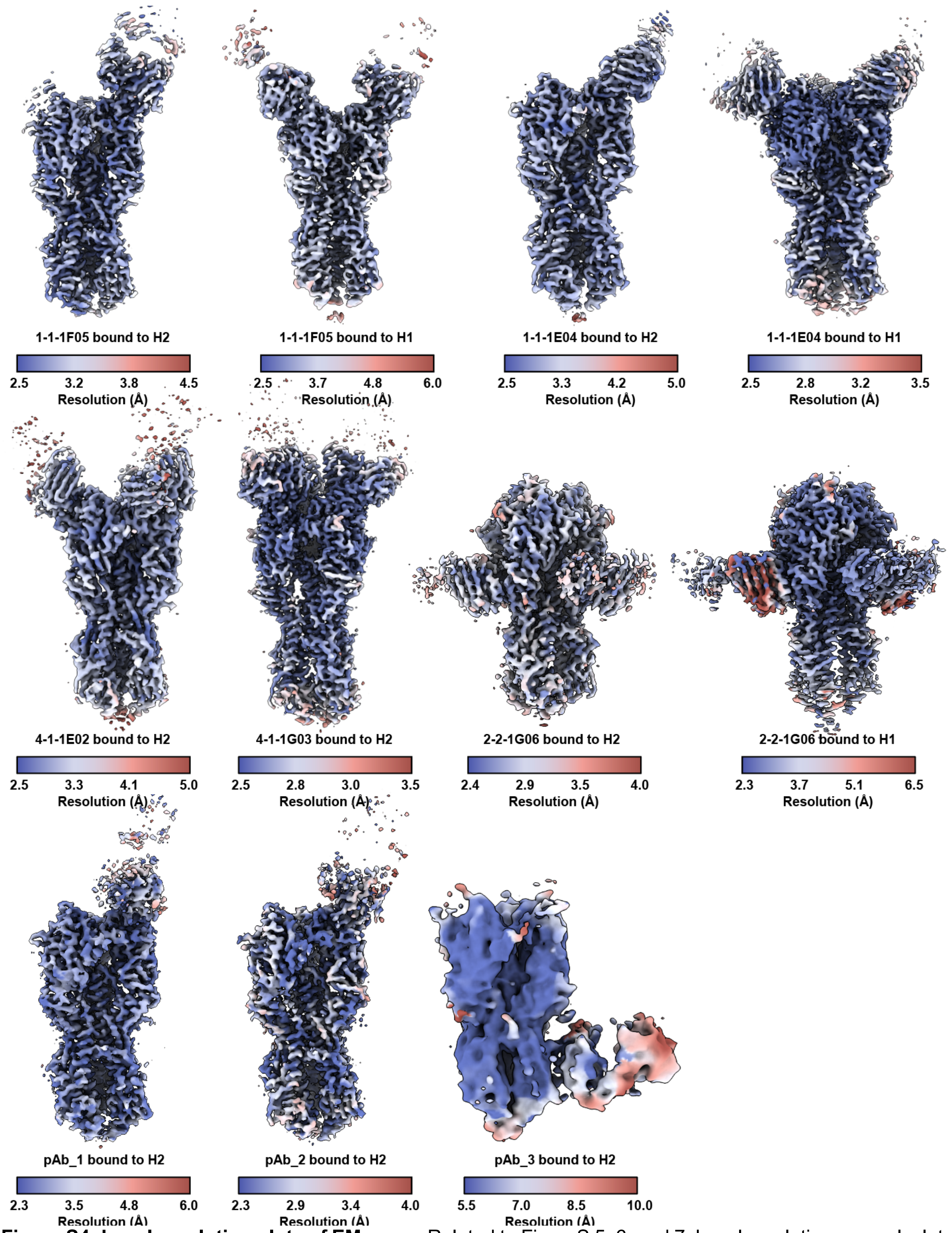


**Figure S4: Local resolution plots of EM maps**. Related to FigureS 5, 6, and 7. Local resolution was calculated according to a 0.143 FSC threshold in cryoSPARC 3.2 and visualized in ChimeraX.

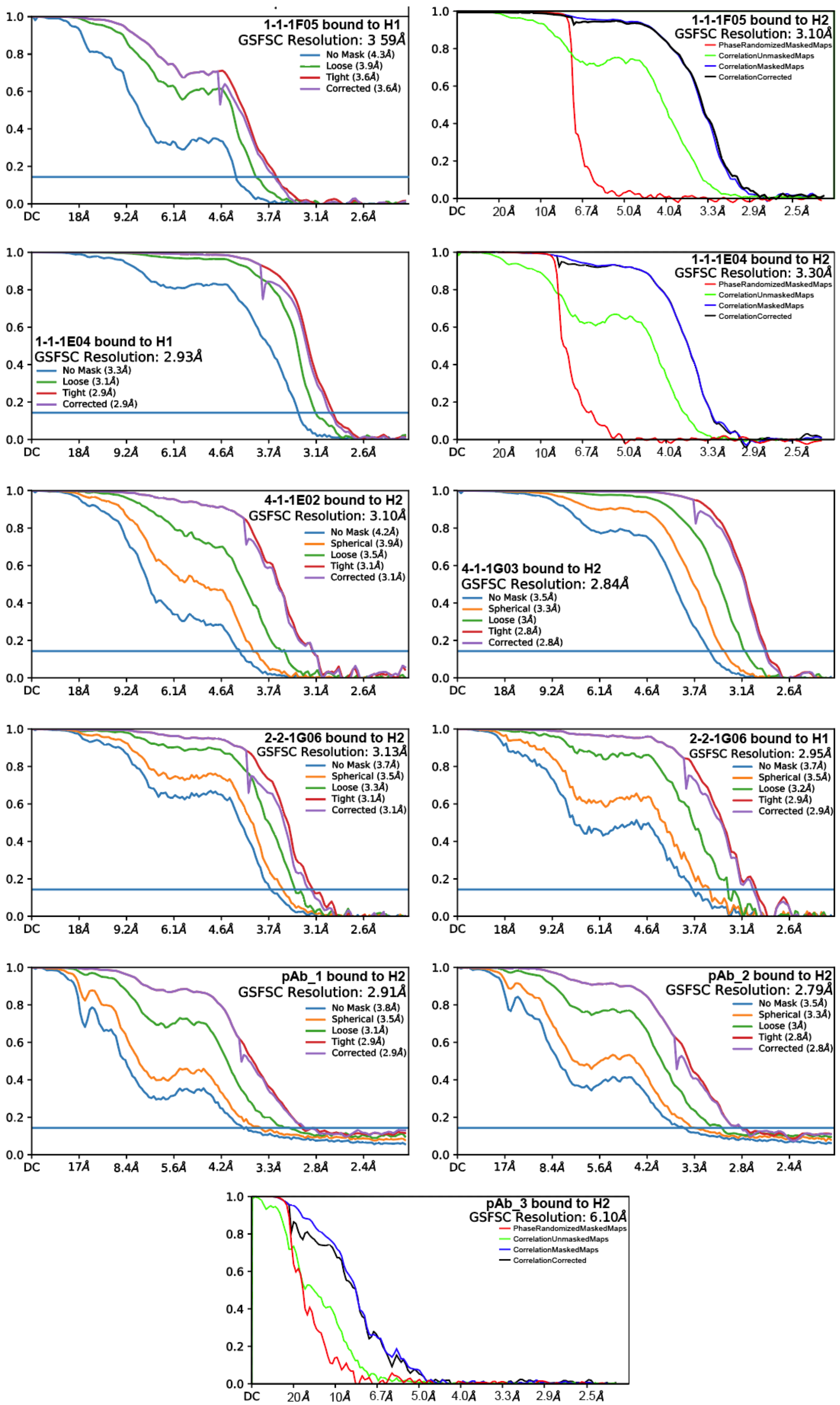


**Figure S5: FSC plots for EM maps.** Related to Figures 5, 6, and 7. Reported resolutions coincide with an FSC cutoff of 0.143. Plots were generated in cryoSPARC 3.2 or Relion.

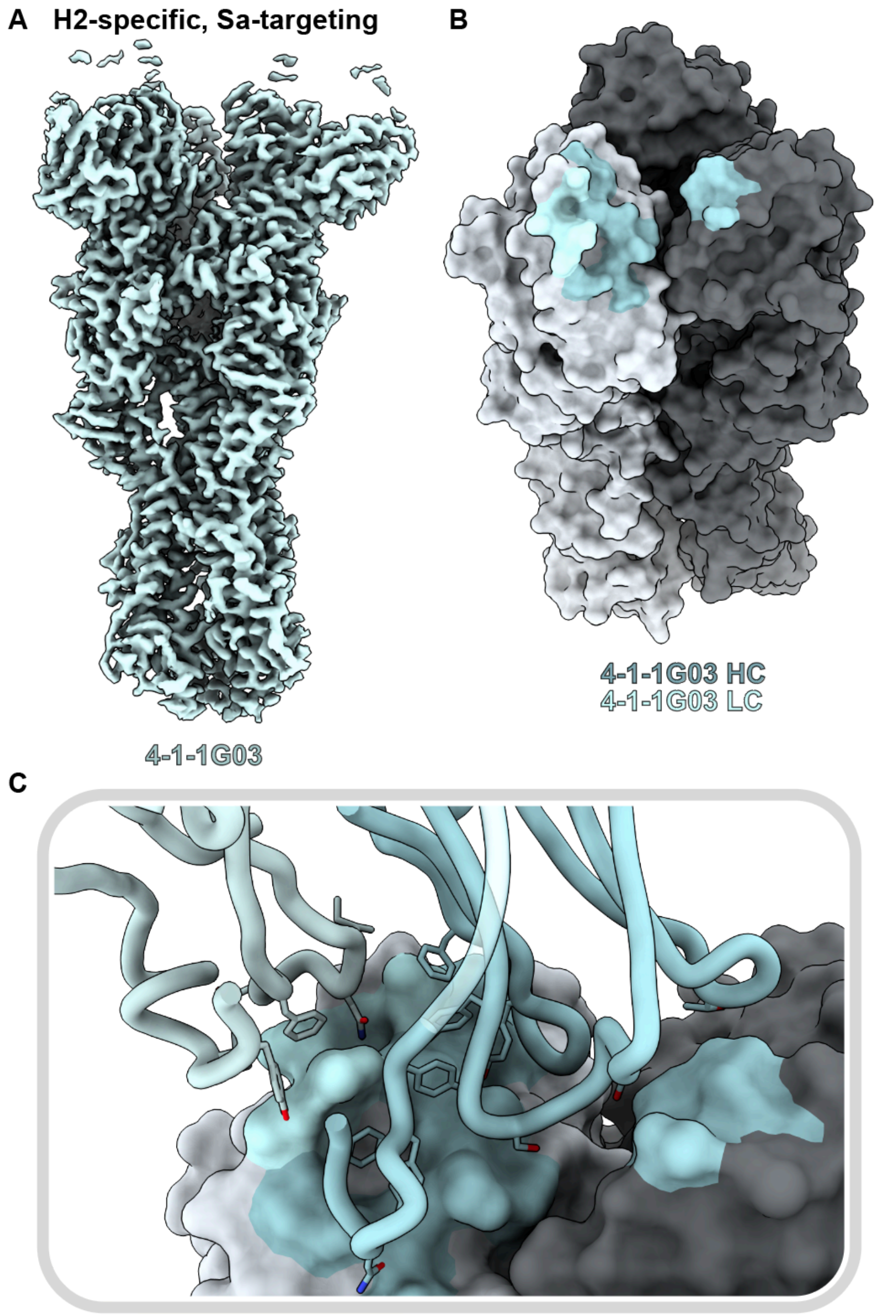


**Figure S6: MAb 4-1-1G03 structural characterization**. Related to Figure 6. (A) cryoEM density map of 4-1-1G03 Fab complexed with H2. (B) Antibody footprint of 4-1-1G03 colored to indicate heavy and light chain interactions on H2. (C) Antibody loop interactions with the SA epitope with key residues shown.

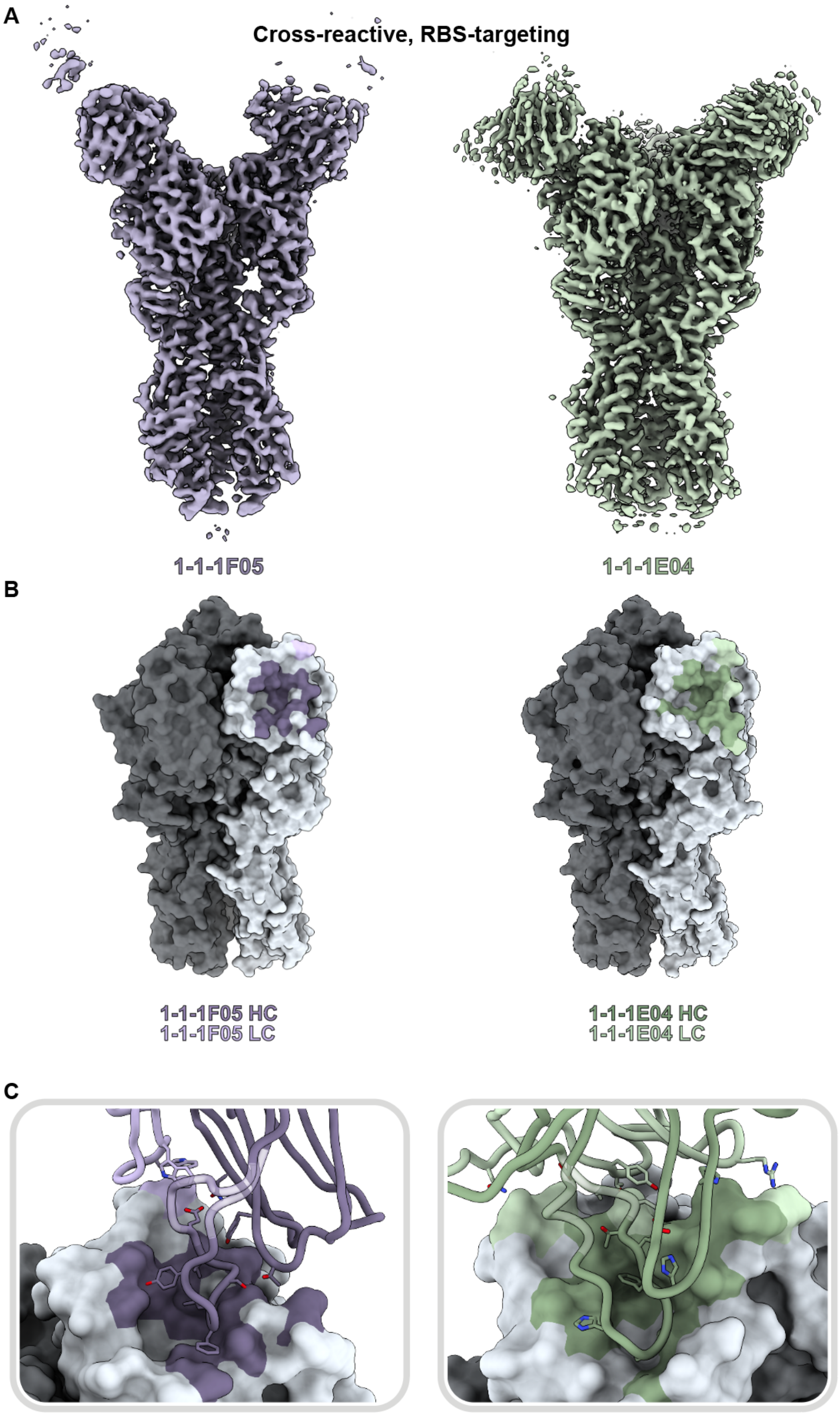


**Figure S7: Cross-reactive mAb binding to H1 NC99.** Related to Figure 6. (A) Cryo-EM reconstructions of mAbs 1-1-1F05 and 1-1-1E04 bound to H1 NC99. Antibody footprint of 1-1-1F05 and 1-1-1E04 colored to indicate heavy and light chain interactions on H1 NC99. (C) Antibody loop interactions with H1 NC99.

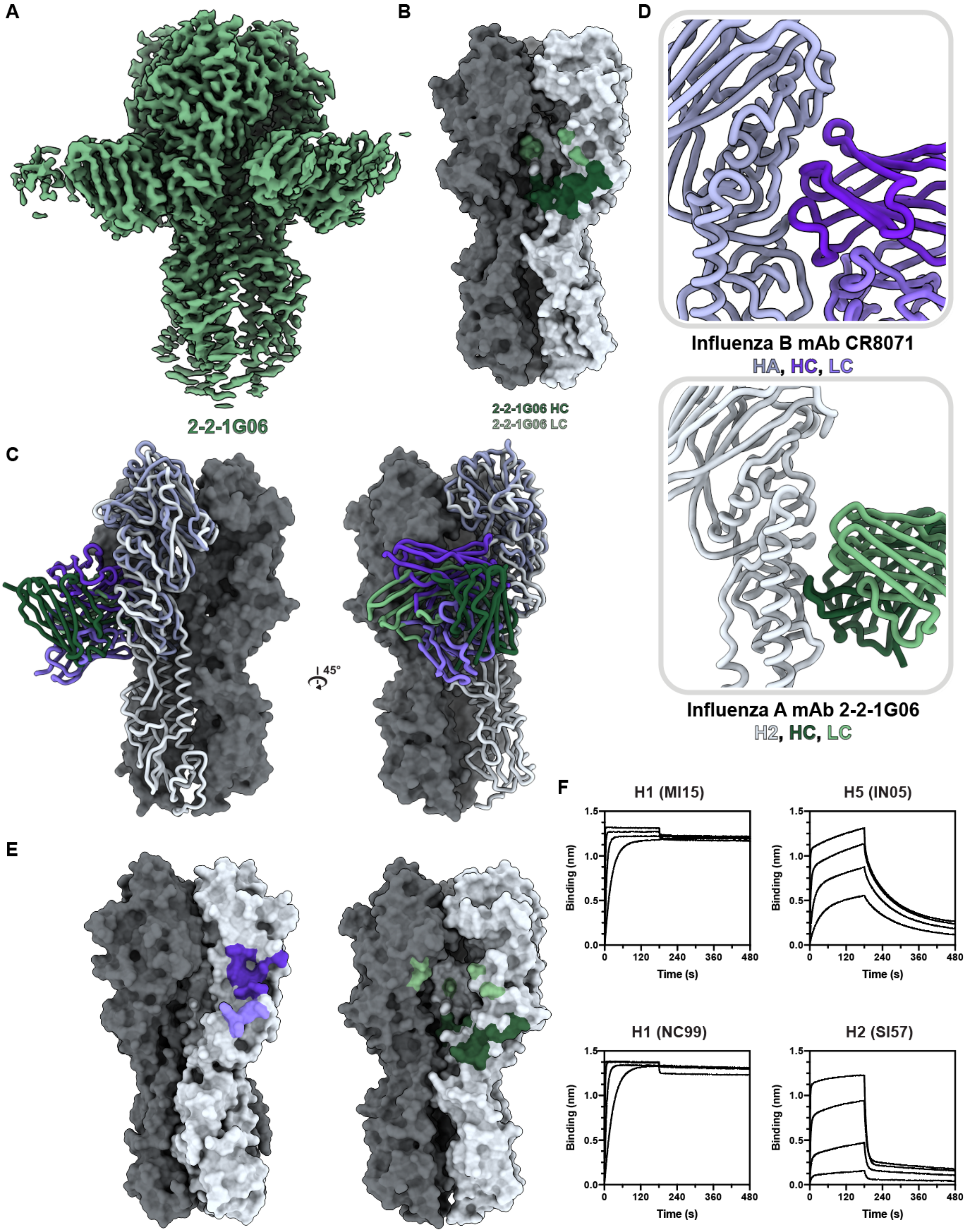

A
2-2-1G06
B
2-2-1G06 HC
2-2-1G06 LC
C
45°
D
Influenza B mAb CR8071
HA, HC, LC
Influenza A mAb 2-2-1G06
H2, HC, LC
E
F
H1 (MI15)
H5 (IN05)
H1 (NC99)
H2 (SI57)
Binding (nm)
Time (s)
0.0
0.5
1.0
1.5
0
120
240
360
480

**Figure S8: MAb 2-2-1G06 interaction with H1 and comparison to CR8071 and binding kinetics**. Related to Figure 7. (A) cryoEM density map of 2-2-1G06 Fab complexed with H1. (B) Antibody footprint of 2-2-1G06 colored to indicate heavy and light chain interactions on H1. (C & D) Comparison of influenza B mAb CR8071 (purple; PDB 4FQJ) bound to HA (PDBs 4FQJ and 4M44) and 2-2-1G06 (green) bound to H2. (E) Antibody footprints of 2-2-1G06 on H2 and CR8071 on HA (PDBs 4FQJ and 4FQM). (F) BLI sensorgrams indicating immobilized mAb 2-2-1G06 binding to H1, H2, or H5 HAs at concentrations of 3200, 800, 200, and 50 nM.

**Table S1: nsEMPEM map and deposition details.** Related to Figures 2 and 4.

| | EMDB ID | Time point | # particles (composite) | Microscope | Pixel size |
|---|---|---|---|---|---|
| Donor 1-1 (H2) | EMD-41514 | 0 | 7,863 | Tecnai TF20 | 1.77 |
| | EMD-41515 | 4 | 11,200 | Tecnai Spirit | 2.06 |
| | EMD-41516 | 16 | 17,496 | Tecnai Spirit | 2.06 |
| | EMD-41517 | 20 | 51,147 | Tecnai Spirit | 2.06 |
| Donor 1-2 (H2) | EMD-41518 | 0 | 2,167 | Tecnai Spirit | 2.06 |
| | EMD-41519 | 4 | 9,223 | Tecnai TF20 | 1.77 |
| | EMD-41520 | 16 | 3,572 | Tecnai Spirit | 2.06 |
| | EMD-41521 | 20 | 57,429 | Tecnai Spirit | 2.06 |
| Donor 1-3 (H2) | EMD-41522 | 20 | 33,782 | Tecnai Spirit | 2.06 |
| Donor 2-1 (H2) | EMD-41523 | 4 | 37,664 | Tecnai TF20 | 1.77 |
| | EMD-41524 | 16 | 80,050 | Tecnai Spirit | 2.06 |
| | EMD-41525 | 20 | 48,917 | Tecnai Spirit | 2.06 |
| Donor 2-2 (H2) | EMD-41526 | 0 | 5,252 | Tecnai TF20 | 1.77 |
| | EMD-41527 | 4 | 23,361 | Tecnai Spirit | 2.06 |
| | EMD-41528 | 16 | 12,935 | Tecnai Spirit | 2.06 |
| | EMD-41529 | 20 | 39,714 | Tecnai Spirit | 2.06 |
| Donor 2-3 (H2) | EMD-41530 | 4 | 19,649 | Tecnai Spirit | 2.06 |
| | EMD-41531 | 16 | 3,437 | Tecnai TF20 | 1.77 |
| | EMD-41532 | 20 | 16,270 | Tecnai Spirit | 2.06 |
| Donor 3-1 (H2) | EMD-41533 | 0 | 7,280 | Tecnai Spirit | 2.06 |
| | EMD-41534 | 4 | 27,785 | Tecnai TF20 | 1.77 |
| | EMD-41535 | 16 | 31,686 | Tecnai Spirit | 2.06 |
| | EMD-41536 | 20 | 33,852 | Tecnai Spirit | 2.06 |
| Donor 3-2 (H2) | EMD-41537 | 0 | 20,800 | Tecnai TF20 | 1.77 |
| | EMD-41538 | 4 | 19,649 | Tecnai Spirit | 2.06 |
| | EMD-41539 | 16 | 41,400 | Tecnai TF20 | 1.77 |
| | EMD-41540 | 20 | 106,050 | Tecnai Spirit | 2.06 |
| Donor 3-3 (H2) | EMD-41541 | 0 | 30,603 | Tecnai Spirit | 2.06 |
| | EMD-41542 | 4 | 27,331 | Tecnai Spirit | 2.06 |
| | EMD-41543 | 16 | 19,700 | Tecnai Spirit | 2.06 |
| | EMD-41544 | 20 | 59,878 | Tecnai Spirit | 2.06 |
| Donor 4-1 (H2) | EMD-41545 | 0 | 27,266 | Tecnai Spirit | 2.06 |
| | EMD-41546 | 4 | 54,296 | Tecnai Spirit | 2.06 |
| | EMD-41547 | 16 | 89,300 | FEI Talos | 1.98 |
| | EMD-41548 | 20 | 93,574 | Tecnai Spirit | 2.06 |
| Donor 4-2 (H2) | EMD-41549 | 0 | 18,505 | Tecnai Spirit | 2.06 |
| | EMD-41550 | 4 | 50,289 | Tecnai Spirit | 2.06 |
| | EMD-41551 | 16 | 10,340 | Tecnai Spirit | 2.06 |
| | EMD-41552 | 20 | 49,813 | Tecnai TF20 | 1.77 |
| Donor 4-3 (H2) | EMD-41553 | 0 | 11,809 | Tecnai Spirit | 2.06 |
| | EMD-41554 | 4 | 82,814 | Tecnai Spirit | 2.06 |
| | EMD-41555 | 16 | 14,356 | Tecnai Spirit | 2.06 |
| | EMD-41556 | 20 | 66,790 | FEI Talos | 1.98 |
| Donor 1-1 (H1) | EMD-41557 | 0 | 33,500 | Tecnai TF20 | 1.77 |
| | EMD-41558 | 4 | 26,600 | Tecnai TF20 | 1.77 |
| | EMD-41559 | 16 | 19,300 | Tecnai TF20 | 1.77 |
| | EMD-41560 | 20 | 53,500 | Tecnai TF20 | 1.77 |
| Donor 2-2 (H1) | EMD-41561 | 0 | 34,000 | Tecnai TF20 | 1.77 |
| | EMD-41562 | 4 | 88,000 | Tecnai TF20 | 1.77 |
| | EMD-41563 | 16 | 65,400 | Tecnai TF20 | 1.77 |
| | EMD-41564 | 20 | 45,700 | Tecnai TF20 | 1.77 |

**Table S2: Cryo-EM map and atomic model refinement.** Related to Figures 5, 6, and 7.

| | 1-1-1F05 bound to H2 | 1-1-1F05 bound to H1 | 1-1-1E04 bound to H2 | 1-1-1E04 bound to H1 | 4-1-1E02 bound to H2 | 4-1-1G03 bound to H2 | 2-2-1G06 bound to H2 | 2-2-1G06 bound to H1 | pAb_1 bound to H2 | pAb_2 bound to H2 | pAb_3 bound to H2 |
|---|---|---|---|---|---|---|---|---|---|---|---|
| **Access codes** | | | | | | | | | | | |
| PDB | 8TP2 | 8TP3 | 8TP4 | 8TP5 | 8TP6 | 8TP7 | 8TP9 | 8TPA | N/A | N/A | N/A |
| EMDB | EMD-41464 | EMD-41465 | EMD-41466 | EMD-41467 | EMD-41468 | EMD-41469 | EMD-41470 | EMD-41471 | EMD-41472 | EMD-41473 | EMD-41474 |
| GenBank | BAF48641.1 | AAP34324.1 | BAF48641.1 | AAP34324.1 | BAF48641.1 | BAF48641.1 | BAF48641.1 | AAP34324.1 | N/A | N/A | N/A |
| **Data collection and processing** | | | | | | | | | | | |
| Microscope | Talos Arctica | Talos Arctica | Talos Arctica | Talos Arctica | Talos Arctica | Talos Arctica | Talos Arctica | Talos Arctica | Titan Krios | Titan Krios | Talos Arctica |
| Magnification | 36,000 | 36,000 | 36,000 | 36,000 | 36,000 | 36,000 | 36,000 | 36,000 | 130,000 | 130,000 | 36,000 |
| Voltage (kV) | 200 | 200 | 200 | 200 | 200 | 200 | 200 | 200 | 300 | 300 | 200 |
| Electron exposure ($e^-/Å^2$) | 53.5 | 49.0 | 49.2 | 46.2 | 49.0 | 49.0 | 46.2 | 48.6 | 49.7 | 49.7 | 50.3 |
| Defocus range (µm) | -0.7 to -2 | -0.7 to -2 | -0.7 to -2 | -0.7 to -2 | -0.7 to -2 | -0.7 to -2 | -0.7 to -2 | -0.7 to -2 | -0.7 to -2 | -0.7 to -2 | -0.7 to -2 |
| Pixel size (Å) | 1.150 | 1.15 | 1.150 | 1.15 | 1.150 | 1.150 | 1.150 | 1.150 | 1.045 | 1.045 | 1.045 |
| Imposed Symmetry | C1 | C1 | C1 | C3 | C3 | C3 | C3 | C3 | C1 | C1 | C1 |
| Final particle number | 230,649 | 105,408 | 117,851 | 167,166 | 103,798 | 271,581 | 164,150 | 124,412 | 39,631 | 61,108 | 17,933 |
| Map resolution (Å) | 3.1 | 3.6 | 3.3 | 2.9 | 3.1 | 2.8 | 3.1 | 3.0 | 2.9 | 2.8 | 6.1 |
| FSC Threshold | 0.143 | 0.143 | 0.143 | 0.143 | 0.143 | 0.143 | 0.143 | 0.143 | 0.143 | 0.143 | 0.143 |
| Map sharpening B-factor ($Å^2$) | -37.9 | -99.7 | -40.8 | -95.0 | -94.9 | -105.9 | -99.6 | -83.7 | -56.3 | -58.8 | -138.4 |
| **Model refinement and validation** | | | | | | | | | | | |
| Total Residues | 1697 | 1720 | 1697 | 2223 | 2153 | 2163 | 2157 | 2172 | | | |
| Amino-acids | 1678 | 1700 | 1678 | 2205 | 2126 | 2145 | 2142 | 2154 | | | |
| Carbohydrates | 19 | 20 | 18 | 18 | 27 | 18 | 15 | 18 | | | |
| RMSD Lengths (Å) | 0.020 | 0.020 | 0.020 | 0.021 | 0.021 | 0.021 | 0.020 | 0.021 | | | |
| RMSD Angles (°) | 1.7 | 1.7 | 1.8 | 1.9 | 1.8 | 1.8 | 1.8 | 1.8 | | | |
| **Ramachandran** | | | | | | | | | | | |
| Outliers (%) | 0 | 0 | 0 | 0 | 0 | 0 | 0 | 0 | | | |
| Allowed (%) | 2.2 | 1.7 | 2.7 | 1.8 | 1.2 | 1.7 | 1.9 | 1.3 | N/A | N/A | N/A |
| Favored (%) | 97.8 | 98.28 | 97.4 | 98.21 | 98.8 | 98.3 | 98.1 | 98.7 | | | |
| Rotamer outliers (%) | 0 | 0.07 | 0 | 0.11 | 0 | 0 | 0 | 0 | | | |
| Clash score | 1.6 | 0.8 | 2.9 | 1.6 | 1.9 | 1.4 | 2.3 | 1.5 | | | |
| Molprobity score | 0.94 | 0.75 | 1.21 | 0.91 | 0.95 | 0.88 | 1.00 | 0.89 | | | |
| FSC model (0/0.143/0.5) | 2.7/2.8/3.1 | 3.5/3.5/3.9 | 2.9/3.0/3.3 | 2.7/2.8/3.1 | 3.0/3.1/3.3 | 2.8/2.8/3.0 | 3.0/3.1/3.3 | 2.9/2.9/3.2 | | | |
| EMRinger score | 4.2 | 2.4 | 4.0 | 3.4 | 3.7 | 4.8 | 3.5 | 4.7 | | | |

**Table S3: nsEM map and deposition details for monoclonal immune complexes.** Related to Figures 4 and 5.

| Monoclonal ID | EMDB ID | Breadth | # particles (composite) | Symmetry | Microscope | Pixel size (Å) |
|---|---|---|---|---|---|---|
| **2-2-1E08** | EMD-41683 | H2 | 3,672 | C1 | Tecnai Spirit | 2.06 |
| **2-2-1C06** | EMD-41684 | H2 | 11,091 | C1 | Tecnai Spirit | 2.06 |
| **4-1-1E02** | EMD-41685 | H2 | 25,927 | C1 | Tecnai Spirit | 2.06 |
| **4-1-1G03** | EMD-41686 | H2 | 14,494 | C3 | Tecnai Spirit | 2.06 |
| **1-3-1F08** | EMD-41687 | H2 | 4,424 | C1 | Tecnai Spirit | 2.06 |
| **1-1-1F05** | EMD-41688 | cross-reactive | 2,741 | C1 | Tecnai Spirit | 2.06 |
| **2-2-1F01** | EMD-41689 | cross-reactive | 22,641 | C1 | Tecnai Spirit | 2.06 |
| **2-2-1G06** | EMD-41690 | cross-reactive | 2,651 | C1 | Tecnai Spirit | 2.06 |
| **1-2-189-34** | EMD-41691 | cross-reactive | 29,435 | C3 | Tecnai F20 | 1.77 |
| **1-1-1A09** | EMD-41692 | cross-reactive | 17020.00 | C3 | Tecnai F20 | 1.77 |
| **1-1-2A11** | EMD-41693 | cross-reactive | 4,231 | C3 | Tecnai F20 | 1.77 |
| **1-1-2E05** | EMD-41694 | cross-reactive | 8,762 | C3 | Tecnai F20 | 1.77 |
